# Landscape restoration strategies differ in their predicted benefits for species persistence

**DOI:** 10.64898/2026.10.06.756954

**Authors:** Robin Bauknecht, Robert J. Fletcher

## Abstract

Global biodiversity recovery demands restoration at landscape scales, where ecological outcomes are shaped not only by habitat amount but also spatial arrangement. Alternative landscape restoration strategies can increase habitat amount, improve quality, and alter configuration in fundamentally different ways and may therefore vary in their efficiency. Here, we conceptualise five landscape restoration strategies and use a spatially explicit metapopulation framework to compare their ability to increase predicted species persistence across heterogeneous landscapes and trait space defined by dispersal ability and specialisation. We find that restoration outcomes are strongly trait-dependent, with restoration of existing degraded habitats yielding the highest benefits – particularly for specialists. Improvement of the landscape matrix to create low-quality habitat primarily benefits generalists, while strategies creating new habitat patches or enlarging patches provide more consistent but modest gains. These results reveal fundamental trade-offs among restoration strategies and highlight the utility of trait-explicit approaches to inform their implementation and evaluation.

## Introduction

Human-driven environmental degradation has dramatically altered ecosystems worldwide. Seventy-five percent of the terrestrial environment is now significantly modified by human activities (IPBES 2019), wildlife populations are declining rapidly across many regions and taxa (Dirzo *et al*. 2014), and global biodiversity indicators show sustained long-term downward trends (Butchart *et al*. 2010). In response to the scale of this degradation, the restoration of damaged ecosystems has gained unprecedented global momentum. The UN Decade on Ecosystem Restoration (2021–2030) has called for a “global movement to ramp up restoration” (UNEP & FAO 2020) and governments and organizations have subsequently pledged to restore hundreds of millions of hectares of degraded land and mobilize billions of dollars in support of these efforts (German Federal Ministry for the Environment & IUCN 2011; United Nations Climate Summit 2014; WWF et al. 2020). This rapid expansion reflects a growing belief that restoration is a powerful tool to halt ecological decline and to support biodiversity recovery at a global scale.

Such global efforts necessarily require habitat restoration at the scale of large landscapes. However, restoration science has traditionally been predominantly site-focused, with comparatively less integration of landscape-scale factors such as connectivity, patch geometry, and the composition of surrounding habitats into restoration design and evaluation (Brudvig 2011; Hobbs & Norton 1996). Yet the factors that ultimately determine restoration success – and failure – may differ fundamentally between local and landscape scales (Levin 1992). While local conditions govern habitat suitability and the extent to which sites meet species’ niche requirements (Soberon & Peterson 2005), landscape-scale properties such as habitat amount, spatial configuration, and connectivity shape dispersal, colonization, and ultimately long-term population persistence (Hanski 1999, 1998). At this scale, restoration becomes inherently geometric: interventions can influence ecological processes not only by altering how much habitat exists or how suitable it is, but also by changing where habitat is located and how it is arranged in space. Although restoration ecology increasingly acknowledges the need to incorporate landscape context, mechanistic understanding of how alternative spatial implementations of restoration translate into persistence outcomes across heterogeneous landscapes remains limited (Brudvig 2011; Gilby *et al*. 2018; von Holle *et al*. 2020; Menz *et al*. 2013; Perring *et al*. 2015).

Metapopulation theory provides a framework for understanding the processes underlying persistence in fragmented landscapes, emphasizing the roles of habitat size, quality, and spatial configuration in shaping colonization-extinction dynamics and regional (metapopulation) persistence (Hanski 1999, 1998). From a metapopulation perspective, restoration interventions can be broadly grouped into a number of distinct mechanisms by which they alter the landscape structure and suitability underlying species persistence. We identify five distinct approaches, what we hereafter collectively term ‘landscape restoration strategies’ (Fig. 1A). Habitat may be added as new, discrete patches that support additional local populations and/or act as ‘stepping stones’ connecting habitats (Fig. 1A I; Gilpin 1980; Saura et al. 2014). Existing patches may be enlarged, increasing local carrying capacity and reducing local extinction risk (Fig. 1A II; Falcy & Estades 2007). Patches may be connected to facilitate movement and dispersal among local populations through the creation of corridors (Fig. 1A III; Damschen et al. 2019). Restoration may also enhance the quality of existing habitat without altering geometry (Fig. 1A IV; Moilanen & Hanski 1998) or modify the surrounding landscape matrix to reduce hostility and increase permeability, thereby enhancing dispersal between patches and, in some cases, providing usable or potentially even habitable space (Fig. 1A V; Fletcher et al. 2024; Yamaura et al. 2022). Collectively, these interventions illustrate that restoration is not merely a question of achieving habitat area targets, but of allocating limited resources among alternative spatial mechanisms that influence interacting colonization and extinction processes (Nicol & Possingham 2010). While each of these different potential interventions have long been acknowledged, it remains unknown how they vary in improving the recovery and persistence of species.

**Figure 1.**
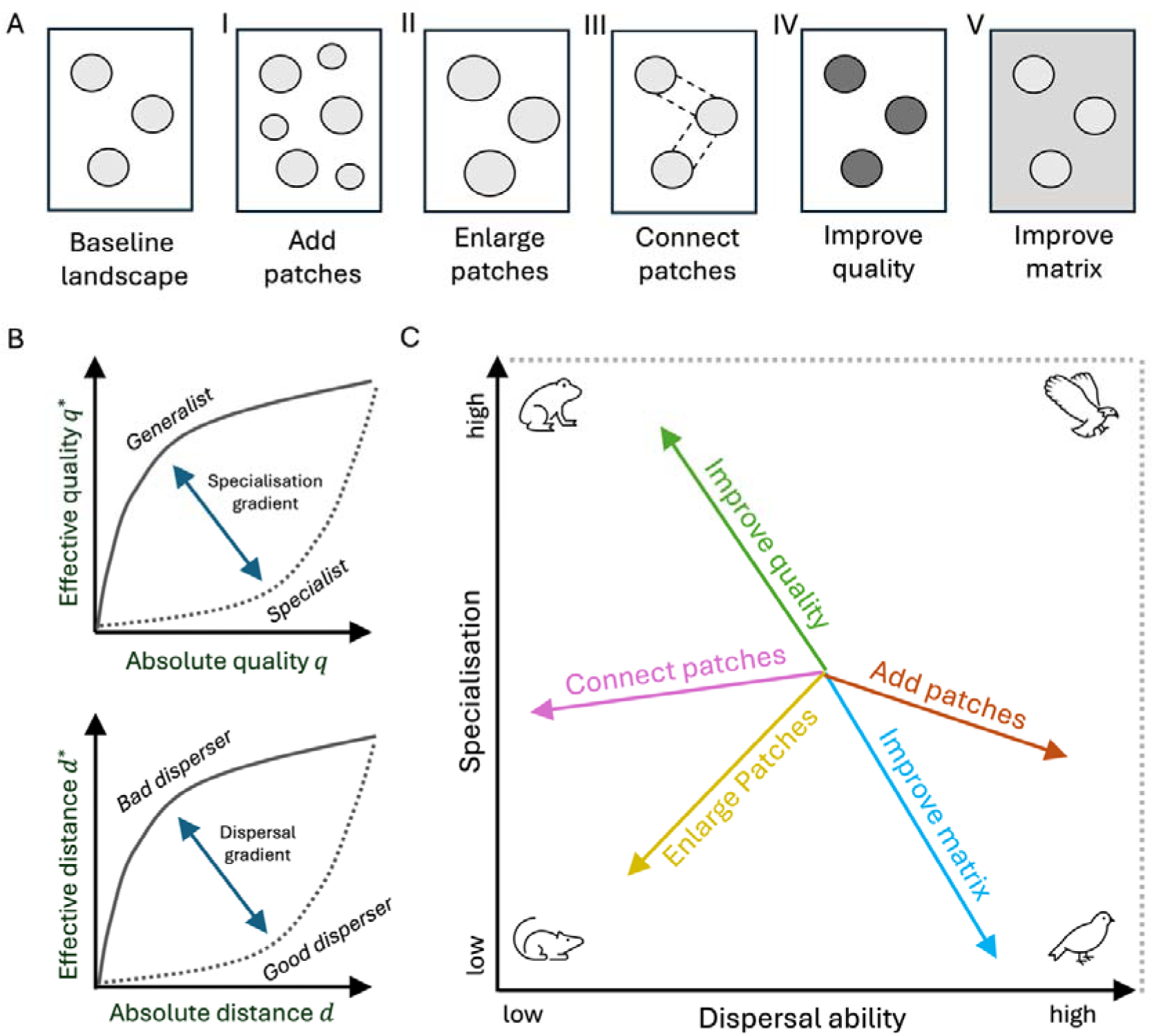
Conceptual overview of landscape restoration strategies and how traits affect their persistence benefit. **(A)** Habitat restoration can be implemented through multiple intervention types that modify either the structure or the quality of a habitat landscape. We consider five broad classes of interventions: (I) adding new habitat patches (“Add patches”), (II) enlarging existing patches (“Enlarge patches”), (III) increasing structural connectivity between patches (“Connect patches”), (IV) improving habitat quality within patches (“Improve quality”), and (V) improving the quality of the surrounding matrix (“Improve matrix”). **(B)** Species differ in how they experience habitat quality and landscape structure depending on their ecological traits. In particular, specialisation determines how absolute habitat quality is translated into effective suitability, whereas dispersal ability determines how spatial distances translate into effective distance. **(C)** As a consequence, the effectiveness of different restoration strategies should depend on species’ positions in trait space defined by dispersal ability and specialisation. Different intervention types are therefore expected to preferentially benefit species with distinct combinations of these traits, leading to systematic variation in restoration efficiency across trait space.

The effects of these landscape restoration strategies on persistence are likely mediated by the species inhabiting the landscape (Hanski et al. 1996). Among the key factors influencing persistence are habitat selection and habitat accessibility (Argote *et al*. 2026; Moilanen & Hanski 1998), both of which are shaped by species traits (Fig. 1B). The specificity in selection preferences can be conceptualized along a specialist–generalist continuum of varying niche breadth (Futuyma & Moreno 1988). Generalist species can exploit a wide range of habitat conditions, whereas specialists depend on specific, high-quality habitats to meet their ecological requirements (Fig. 1B I). Accessibility, in turn, is governed by the interplay of landscape structure with movement and dispersal traits (Brodie et al. 2025; Fahrig 2007). Species with high dispersal ability can potentially traverse fragmented landscapes at relatively low costs, while poor dispersers may be strongly constrained by habitat connectivity (Bonte et al. 2012). Together, specialisation and dispersal ability thus largely determine the effective landscape experienced by a species (Fig 1B). Because these trait axes jointly determine species’ landscapes perception, we expect the efficiency of different restoration strategies to vary systematically across this trait space (Fig. 1C). For example, improving habitat quality may disproportionately benefit specialist species that require high-quality conditions, whereas generalists may respond more strongly to interventions that increase habitat amount or connectivity. However, the direction, magnitude, and generality of this variation in efficiency remain poorly understood, particularly at landscape scales relevant to contemporary restoration initiatives (Brudvig 2011). Resolving these relationships is critical for allocating limited restoration resources in ways that maximize species – and ultimately biodiversity – persistence across diverse species assemblages.

Here, we use a spatially explicit metapopulation framework to examine the persistence benefits of different landscape restoration strategies (Fig. 1). We first ask whether landscape restoration strategies differ in their benefits for increasing species persistence. Second, we ask whether these outcomes vary with species traits. Finally, we ask whether this generates emergent trade-offs when restoring habitat in ecological communities with varying traits. To do so, we simulate a variety of heterogeneous landscapes, implement five broad classes of landscape restoration strategies that modify habitat amount, quality, and spatial structure, and evaluate their effects on species persistence across gradients of dispersal ability and specialisation. Persistence outcomes are quantified using metapopulation capacity (Hanski & Ovaskainen 2000), a mechanistic metric linking landscape structure to long-term persistence potential. By explicitly linking restoration configuration to species traits, our framework provides a general, trait-based perspective on landscape-scale restoration efficiency and highlights the conditions under which different restoration strategies are likely to succeed, or fail, across ecological communities.

## Methods

### Simulating spatially heterogeneous landscapes

We simulated spatially explicit landscapes on a regular 100×100 cell grid. Each cell represents a unit area and is assigned a continuous habitat quality value *q_i∈[0,1]* Cells with *q_i_ = 0* are non-habitat matrix; cells with *q_i_ > 0* are habitat. Habitat patches are defined implicitly as spatially contiguous clusters of habitat cells (4-neighbour connectivity).

We separated two scales of spatial heterogeneity: (i) heterogeneity in the arrangement of habitat patches (hereafter “landscape heterogeneity”) and (ii) heterogeneity in habitat quality within patches (hereafter “habitat heterogeneity”; S1 Fig. S2). Landscape heterogeneity was generated by drawing a Gaussian noise field, smoothing it with a Gaussian kernel, and thresholding the smoothed field to select a fixed number of habitat cells (S1 Fig. S1). The kernel width *σ_L_* controls spatial autocorrelation with larger values producing smoother, coarser patch structure.

Habitat heterogeneity was generated independently from a second, spatially autocorrelated latent quality field (hereafter the “quality potential”), smoothed using a Gaussian kernel with width *σ_q_*. For each landscape, the quality potential was mapped onto habitat cells only; matrix cells retained *q_i_ = 0*. Habitat quality values were monotonically mapped into a bounded interval (*q_i_ ϵ [0.1,1]*) and rescaled so that total effective habitat area *∑_i_q_i_* was constant across landscapes and equal to the number of habitat cells times the mean of the quality range. This ensured that differences among landscapes reflected spatial structure and heterogeneity rather than total habitat quality.

We varied total habitat amount (three levels of 500, 1500, and 2500 habitat cells corresponding to 5%, 15%, and 25% of the total landscape, respectively), landscape heterogeneity (three levels, σ_L_ = 1; 2.5; 5; S1 Fig. S2), and habitat heterogeneity (three levels, *σ_q_ = 1; 2.5; 5*; S1 Fig. S2) in a fully crossed design (27 total combinations). For each combination, we generated 10 independent replicate landscapes, differing only in the random realization of the underlying fields (270 total landscapes).

### Implementing habitat restoration interventions

We implemented five classes of restoration interventions that modify habitat configuration and/or quality whilst controlling for restoration effort (Fig. 1A). In the main analysis, restoration effort was defined as the increase in effective habitat area, *Δ∑i_q__i* from landscape restoration strategies. All interventions were therefore constrained to produce the same increase in effective habitat area. We evaluated three levels of effective habitat area increase *Δ∑iqi*, in {250, 500, 750} corresponding to restoring 2.5%, 5%, and 7.5% of the total landscape from non-habitat to maximum-quality habitat, respectively. We then quantified restoration efficiency as the signed percentage change in metapopulation capacity per unit restoration effort (*%Δλ* / effective habitat area restored). When interventions created new habitat cells, we imposed a minimum quality of *q_min_ = 0.1* to ensure a nontrivial cost per cell and to prevent solutions that allocate infinitesimal quality across many cells. Quality for newly restored cells was derived from the latent quality potential at those locations. Potential values were mapped monotonically into (*q_i_ ϵ [0.1, 1]*), then adjusted to match the specified habitat increase while respecting bounds.

The five intervention types were (Fig. 1A):

1. Add habitat patches. New discrete patches were created in the matrix by randomly selecting and placing patch shapes from an empirical library of patch geometries extracted from replicate landscapes with the same level of spatial heterogeneity. Patch placement was random but constrained to avoid overlap or direct contact with existing habitat. New cells received quality based on their latent potential.
2. Enlarge habitat patches. Existing patches expanded outward into adjacent matrix cells. Expansion proceeded radially from patch boundaries, with each layer defined as the set of matrix cells at equal 4-neighbour distance from the habitat edge. All patches were expanded through one layer before any patch was expanded further, ensuring balanced growth among patches. Newly converted cells received quality based on their latent potential.
3. Connect habitat patches. Disconnected habitat components were linked by adding new habitat along shortest paths between patch boundaries, effectively merging patches through linear expansions. Restoration continued by widening these connections and reinforcing the resulting network until the specified restoration effort was achieved. Newly created cells received quality based on their latent potential. Although this intervention increased habitat area, it did so in a spatially targeted manner that prioritised reducing separation among patches (unlike the “Enlarge habitat patches” strategy).
4. Improve habitat quality. Quality was increased uniformly across existing habitat cells, without altering habitat amount or configuration, until either the specified restoration effort was achieved or cells reached a maximum quality. In landscapes with little initial habitat and high restoration effort, all existing habitat cells could reach maximum quality before the specified increase in effective habitat area was achieved. In such cases, restoration continued by converting adjacent matrix cells. We evaluated scenarios in which this spillover occurred in a separate sensitivity analysis to confirm that it did not drive the main results (S1 Fig. S8).
5. Improve matrix quality. The specified effort was distributed uniformly across matrix cells, effectively converting the matrix into low-quality habitat evenly across the landscape rather than in localized patches. Allocations were not subject to a minimum per-cell threshold.

Detailed descriptions of the algorithmic implementation of each intervention are provided in S1 Text S1. We additionally considered an alternative scenario in which restoration effort was standardised by total area restored, measured as the number of cells restored to maximum habitat quality (*q_i_ = 1*). This scenario represents restoration efforts that maximize habitat quality at locations, in contrast to the primary scenario, in which a common effective habitat area increase can be distributed incrementally across cells. Details and associated results are provided in the Supplementary Information S2.

### Measuring persistence via metapopulation capacity

We quantified the landscapes’ long-term ability to sustain species persistence using metapopulation capacity, defined as the dominant eigenvalue *λ_M_* of a connectivity matrix *M* that captures how spatial structure, habitat amount, and dispersal jointly influence species persistence (Hanski & Ovaskainen 2000). Larger *λ_M_* indicates landscapes better able to support persistent metapopulations. In our pixel-based implementation, each habitat cell is an occupiable unit with quality *q_i_*, building on early work demonstrating that metapopulation processes depend not only on habitat configuration but also on spatial variation in habitat quality (Fleishman *et al*. 2002; Moilanen & Hanski 1998). Connectivity between cells *i* and *j* decays exponentially with their Euclidean distance *d_i,j_* as

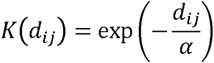

where *α* controls dispersal ability, and larger *α* values correspond to longer-ranged dispersal. A cell’s contribution to colonization scales with its quality, yielding the matrix

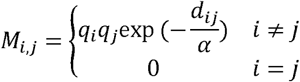

and metapopulation capacity *λ_M_* as the dominant eigenvalue of *M*. Unlike classic patch-based formulations, patch area does not enter explicitly; patch size and habitat amount emerge from the number, contiguity, and quality of habitat cells.

We computed *λ_M_* via power iteration combined with FFT-based convolution (Pavel & David 2013). For computational efficiency, the dispersal kernel was truncated at 100 cells, excluding only the longest diagonal distances. To avoid circular convolution artefacts inherent to FFTs, we implemented zero-padding (i.e., assuming no habitat beyond the landscape boundary), eliminating artificial wrap-around connectivity (Pavel & David 2013). For each replicate landscape, metapopulation capacity was computed for the baseline and every restoration outcome.

While metapopulation capacity provides a mechanistic metric of persistence potential near extinction which is useful for capturing spatially explicit variation in landscape structure (Hanski & Ovaskainen 2000), stochastic patch occupancy models (SPOMs) offer a complementary framework to investigate the underlying colonization-extinction dynamics and the potential for short-term lag effects explicitly. We therefore implemented SPOM simulations to determine if results broadly agree with metapopulation capacity results (Supplementary Information S3).

### Incorporating Dispersal Ability and Specialisation

We represented species traits along two axes: dispersal ability and specialisation. Dispersal ability determines how strongly spatial separation constrains colonization among habitat locations. In fragmented landscapes, species with limited dispersal experience stronger effective isolation among habitat patches, whereas highly mobile species can traverse larger distances and maintain connectivity across the landscape (Bonte *et al*. 2012; Hanski 1999). Dispersal ability was therefore controlled by the kernel scale *α*,which determines the decay of colonization with distance. Specifically, *α* represents the distance at which the dispersal kernel declines to *e_-1_*, or approximately 37%, of its value at zero distance. Smaller *α* values therefore represent short-range dispersers, whereas larger values represent long-range dispersers (Fig. 2B). We evaluated a logarithmically spaced range of 10 *α* values from 1 to 100 pixels, corresponding to dispersal scales ranging from 0.01 to 1 times the linear extent of the 100 × 100-pixel landscapes.

**Figure 2.**
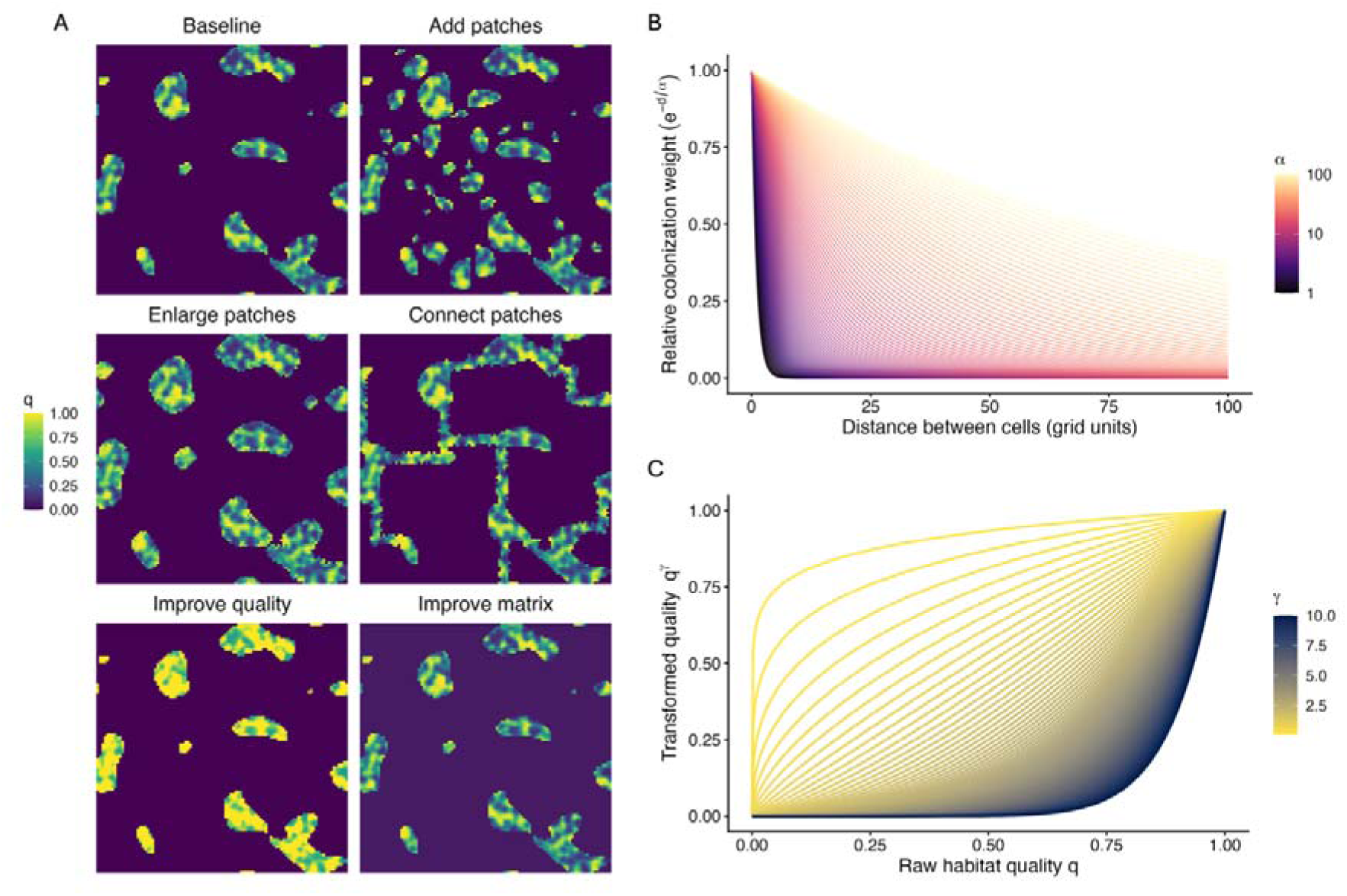
Simulation framework for evaluating restoration interventions across trait space. **(A)** A variety of landscapes were simulated as 100 × 100 grids of habitat quality values *q_i_*, where each cell corresponds to habitat of unit area. Shown here is an example baseline landscape (*∑_i_q_i_ =1500, σ_L_=5,σ_q_=1*) and the associated implementation of the five landscape restoration strategies considered in the analysis (here controlled for effective habitat area increase of *Δ∑_i_q_i_* =500). Landscape and habitat heterogeneity in baseline landscapes is represented in S1 Fig. S2, all baseline landscapes used in the analysis can be found in the compiled code. **(B)** Dispersal ability was represented through a distance-decay colonization, where the scale parameter α controls how rapidly connectivity between habitat declines with distance. Low α values correspond to short-range dispersers that primarily interact locally, whereas high α values generate broader connectivity and increase long-distance colonization contributions. Dispersal kernels were used to compute distance-weighted connectivity among all habitat cells when estimating metapopulation capacity. **(C)** Specialisation was represented using a power transformation of habitat quality values prior to computing metapopulation capacity. Low *γ* values compress quality differences, representing generalists that perceive low- and high-quality habitat more similarly, whereas high *γ* values accentuate quality differences, representing specialists that disproportionately depend on high-quality habitat. A spatial representation of the application of *γ* to landscapes is given in S1 Fig. S3.

Specialist species typically depend on a narrow range of environmental conditions, whereas generalists can exploit a broader range of habitats (Futuyma & Moreno 1988). Here, we represented habitat quality as the degree to which local conditions match a species’ ecological requirements and modelled specialisation by transforming habitat quality using a power function:

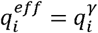

where *γ* controls the species’ sensitivity to habitat quality. This formulation follows approaches that incorporate continuous habitat quality into connectivity-based, landscape metrics (Beyer *et al*. 2020), but here the exponent is interpreted as a species-specific specialisation. Values *γ < 1* compress differences among habitat qualities, representing generalists, whereas *γ > 1* accentuates differences and increases the contribution of high-quality habitat, representing specialists (Fig. 2C). We evaluated 10 logarithmically spaced *γ* values spanning broad generalists (*γ = 0.1*) to strong specialists (*γ* = 10). For each landscape and restoration outcome, we evaluated metapopulation capacity across the fully crossed grid of dispersal (*α*) and specialisation (*γ*) values.

## Results

### Landscape restoration strategies differ in their benefits for species persistence

We applied all five landscape restoration strategies (Fig. 1A I-V) across the full set of simulated landscapes and evaluated outcomes across the dispersal-specialisation trait axes using metapopulation capacity as a proxy for long-term species persistence potential. As expected, across scenarios, restoration increased metapopulation capacity regardless of strategy choice (Fig. 3B). However, the efficiency of the different strategies in doing so varied substantially (Fig. 3A&B). Improving habitat quality had the overall highest median restoration efficiency (Fig. 3B) and was the most frequent optimal strategy, defined here as the intervention yielding the highest restoration efficiency within a given landscape–trait combination, leading to the highest restoration efficiency in approximately *68.9%* of all landscape-trait combinations (Fig. 3A). Improving matrix quality was the second most frequent optimal strategy, yielding the highest restoration efficiency in approximately *26.5%* of cases (Fig. 3A), while the remaining three strategies – adding patches, enlarging patches, and connecting patches – were rarely the single best-performing interventions (< 5%, Fig. 3A). Although improving habitat quality had the highest median restoration efficiency, it also had high variability in efficiency, indicating that its efficiency was highly landscape- and trait-dependent (Fig. 3B). Improving matrix quality similarly exhibited variable performance, with a low median restoration efficiency but occasional high-efficiency outcomes (Fig. 3B). In contrast, adding, enlarging, and connecting patches yielded lower median restoration efficiency, but showed less variability in efficiency (Fig. 3B). The relative patterns among restoration strategies were qualitatively consistent across landscape and habitat heterogeneity and restoration-effort levels, although proportional gains tended to be greater in landscapes with lower initial habitat amount (Figs. S4–S7). The broad relative patterns among strategies were also qualitatively reflected in a complementary stochastic patch occupancy analysis (S3 Fig. S1).

**Figure 3.**
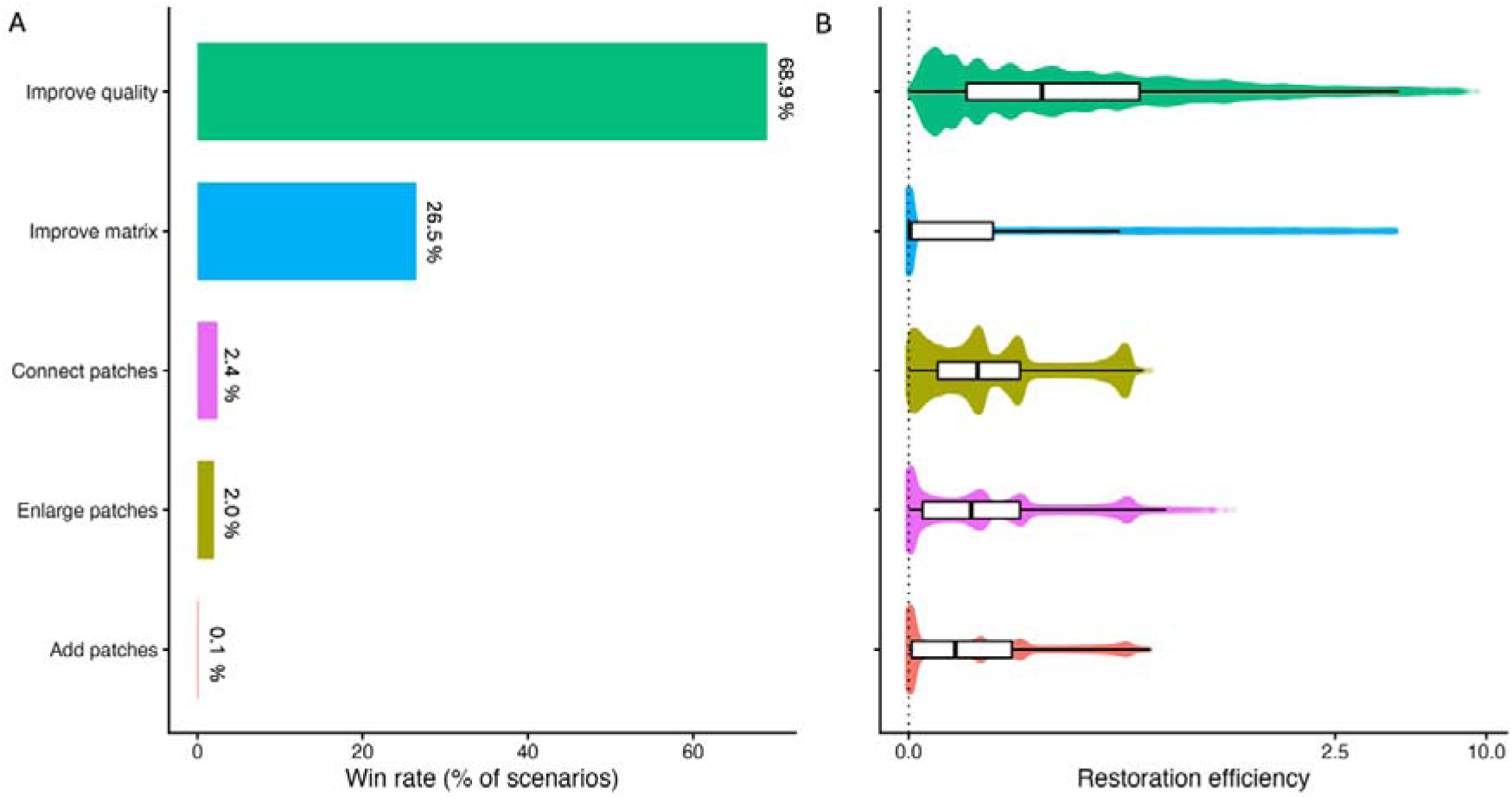
Average effects of landscape restoration strategies on persistence. **(A)** Win rate of each strategy, defined as the proportion of all evaluated scenarios in which the strategy produced the highest restoration efficiency (signed percentage change in metapopulation capacity per unit of effective habitat area restored). Scenarios span the full factorial set of simulated landscapes (habitat amount, landscape heterogeneity, habitat heterogeneity, and replicate) and the full dispersal–specialisation trait grid. Improving habitat quality achieved the highest win rate, matrix improvement was the second most frequent winner, and adding, enlarging, or connecting patches were rarely the most efficient strategy. **(B)** Distribution of restoration efficiency across all landscapes and the dispersal–specialisation trait grid. Violin plots show kernel density; boxes indicate the interquartile range, and the central line is the median. The dotted vertical line marks zero effect. Improving habitat quality yields the highest overall restoration efficiency but also a large spread. Matrix improvement shows the broadest spread, with outcomes ranging from strongly positive to slightly negative. Adding, enlarging, or connecting patches show lower overall effectiveness but less variation. The multimodal “bumps” for add/enlarge/connect patches reflect the variation in efficiency in the three levels of total habitat amount (S1 Fig. S6).

### Species traits mediate the benefits of landscape restoration strategies

The predicted persistence benefits of landscape restoration strategies were strongly trait dependent. Restoration efficiency generally increased with dispersal ability, but the strength of this increase differed among strategies and interacted strongly with specialisation (Fig. 4–6). For adding patches, enlarging patches, and connecting patches, restoration efficiency increased with dispersal ability but varied comparatively little across specialisation classes, producing broadly similar benefits for generalists, intermediate species, and specialists (Fig. 4B; Fig. 5B). In contrast, habitat quality and matrix improvement showed strong but opposing responses to specialisation. Habitat quality improvement became increasingly efficient for specialist species, especially at higher dispersal abilities, whereas matrix improvement was most efficient for generalists and declined sharply with increasing niche specialization (Fig. 4B; Fig. 5B).

**Figure 4.**
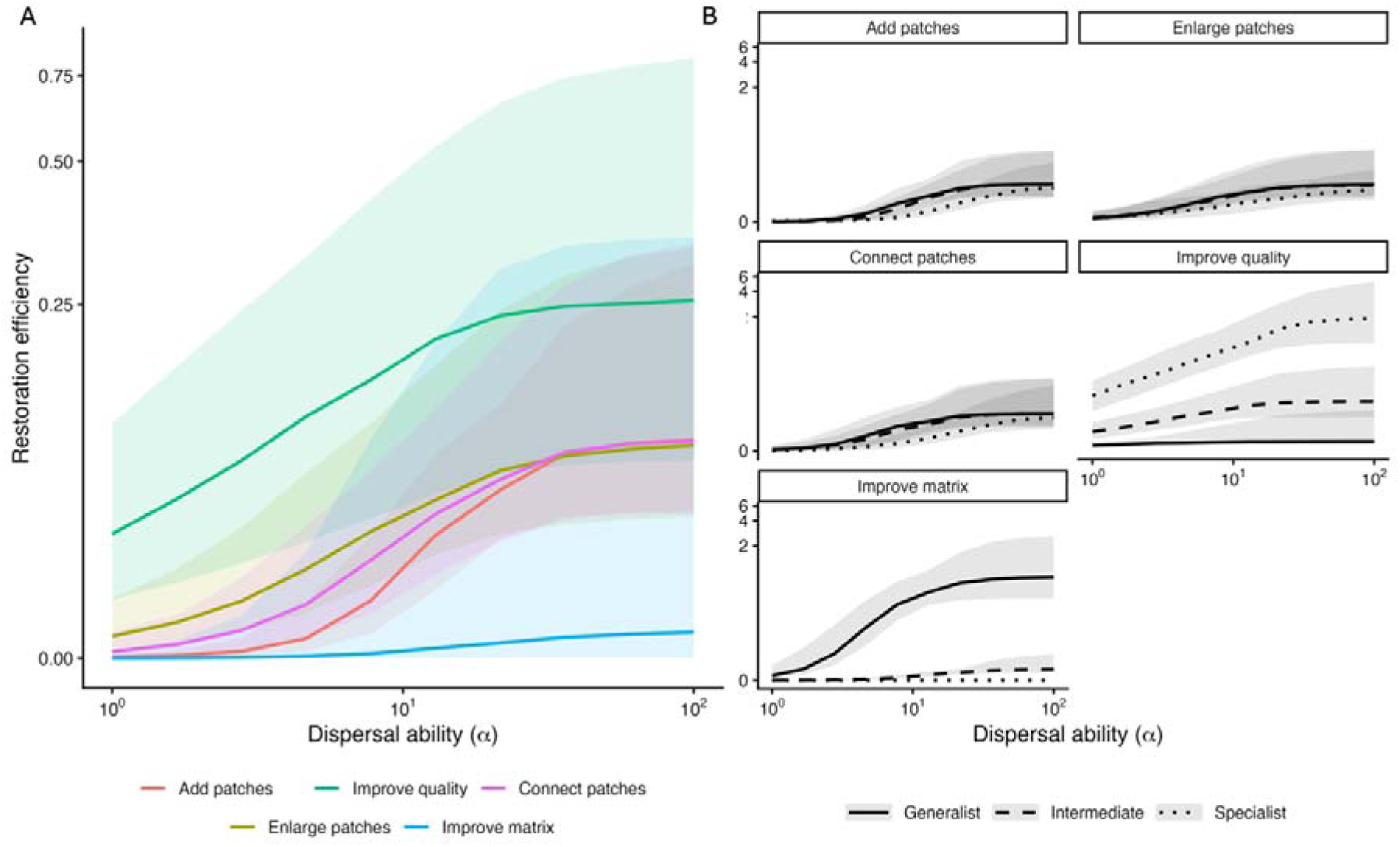
The persistence benefits of restoration strategies across dispersal ability. **(A)** Restoration efficiency (signed percentage change in metapopulation capacity per unit effective habitat area restored) across dispersal ability (α) for each restoration intervention. Lines show median restoration efficiency across all simulated landscapes and specialisation values (effects of individual factors are represented in S1 Fig. S4-7), with shaded bands indicating the interquartile range. Restoration efficiency generally increased with dispersal ability, but interventions differed strongly in magnitude: improving habitat quality yielded the highest efficiency across most of the dispersal gradient, whereas improving the matrix produced consistently low efficiency. Adding patches, enlarging patches, and connecting patches showed intermediate and broadly similar efficiencies. **(B)** Efficiency–dispersal relationships shown for representative generalist, intermediate, and specialist species (lowest, middle, and highest values of the simulated *γ* range). The dominance of habitat quality improvement was driven primarily by strong gains for specialist species, whereas matrix improvement disproportionately benefitted generalists and was ineffective for specialists. Habitat-adding interventions showed comparatively weak differentiation across specialisation classes.

**Figure 5.**
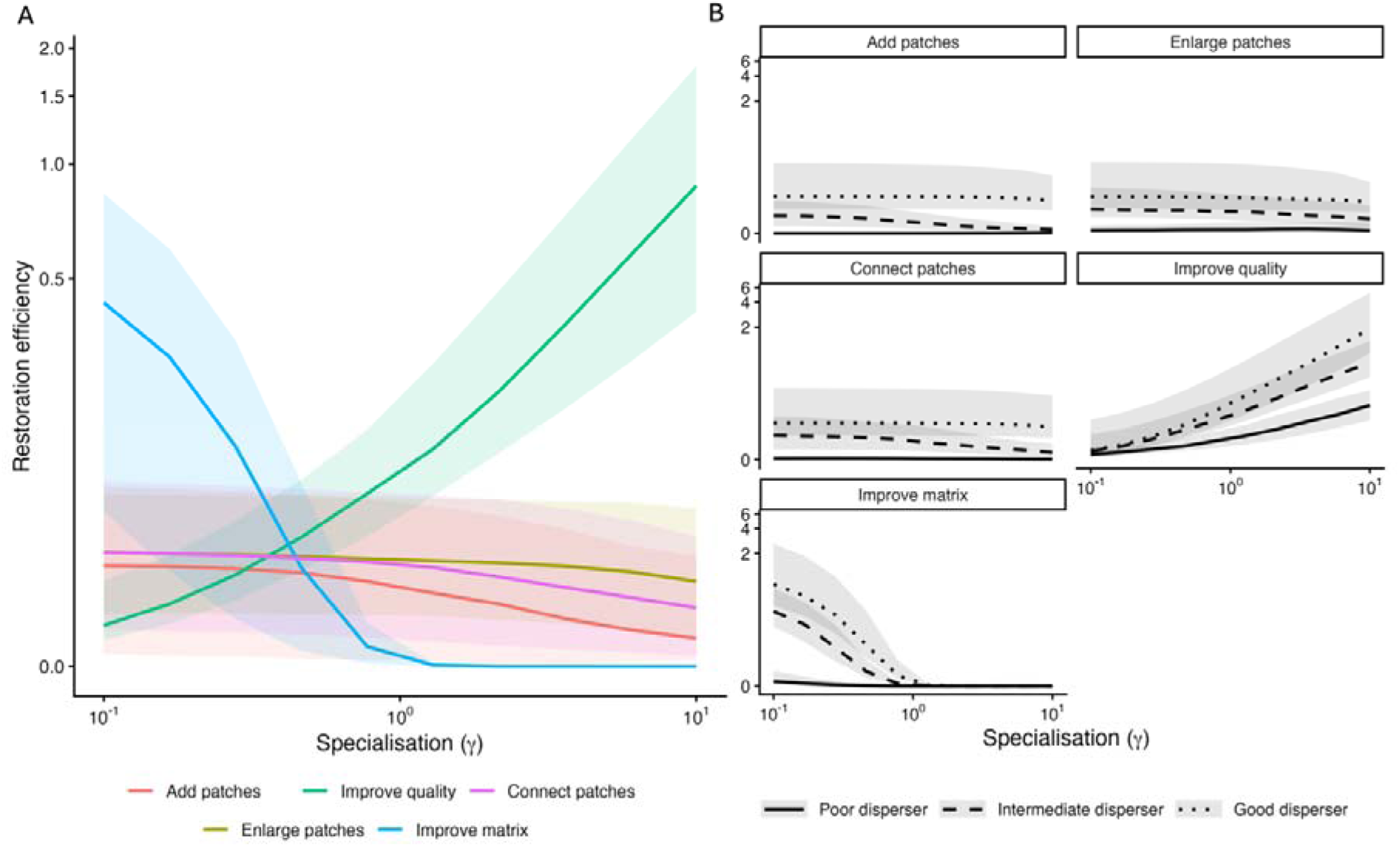
The persistence benefits of restoration strategies across specialisation. **(A)** Restoration efficiency (signed percentage change in metapopulation capacity per unit effective habitat area restored) across specialisation (*γ*) for each restoration intervention. Lines show median restoration efficiency across all simulated landscapes and dispersal ability values (effects of individual factors are represented in S1 Fig. S4-7), with shaded bands indicating the interquartile range. Restoration efficiency varied strongly with specialisation for interventions affecting habitat quality and matrix permeability: improving habitat quality increased sharply with*γ,* yielding highest efficiency for specialists, whereas improving the matrix declined steeply with *γ* and approached zero for specialist species. In contrast, adding patches, enlarging patches, and connecting patches showed comparatively weak dependence on specialisation, with broadly stable efficiency across *γ.* **(B)** Efficiency-specialisation relationships shown for representative low, intermediate, and high dispersal abilities (lowest, middle, and highest values of the simulated α range). The positive *γ*-dependence of habitat quality improvement was strongest for good dispersers, for which restoration efficiency increased most steeply toward high *γ*. Conversely, matrix improvement became rapidly ineffective with increasing *γ* across dispersal classes. For the three habitat-adding interventions, dispersal ability primarily shifted overall efficiency magnitude but did not qualitatively alter their weak *γ*-dependence.

**Figure 6.**
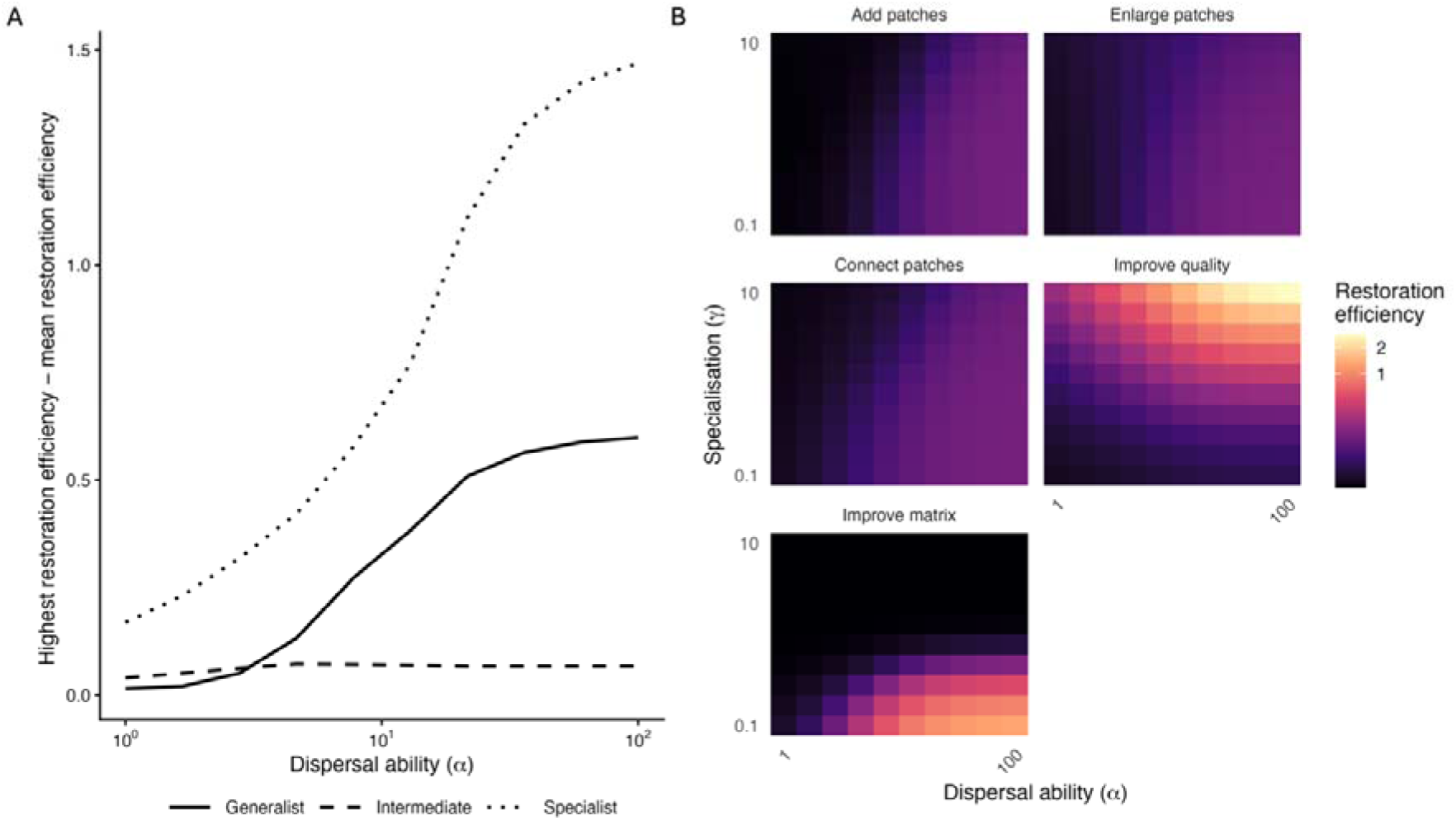
Restoration strategies show strong efficiency trade-offs for mobile species. **(A)** Difference between the most efficiently restoration strategy and the mean (highest restoration efficiency − mean restoration efficiency) across dispersal ability (α), shown for representative low, intermediate, and high specialisation values corresponding to the lowest, middle, and highest values of the simulated *γ* range. Restoration efficiency is defined as the signed percentage change in metapopulation capacity per unit effective habitat area restored. Differences among strategies increased with dispersal ability and were largest for species at the extremes of the specialisation spectrum (generalists and specialists), whereas intermediate specialisation species showed comparatively small variation in achievable restoration efficiency across strategies. At low dispersal ability, all strategies yielded similarly low efficiency, resulting in limited differences between best and average performance. **(B)** Mean restoration efficiency across the full dispersal–specialisation trait grid for each restoration intervention. Heatmaps illustrate that the strong divergence in panel A is primarily driven by opposing trait-dependent responses to improving habitat quality versus improving the matrix: habitat quality improvements yielded high efficiency for specialist species, particularly at high α, whereas matrix improvements produced high efficiency primarily for broad generalists. In contrast, patch-based interventions (adding, enlarging, connecting patches) showed intermediate efficiency and comparatively weak variation across trait space. Together, these results indicate that for highly mobile species, restoration planning may involve a trade-off between interventions that preferentially promote specialist persistence versus those that benefit generalists.

These opposing trait responses generated trade-offs in restoration efficiency across trait space (Fig. 6A–B). Differences among strategies increased strongly with dispersal ability and were greatest at the extremes of the specialisation spectrum. For highly mobile species, habitat quality improvement produced the largest persistence gains for specialists, whereas matrix improvement produced the largest gains for generalists. In contrast, adding, enlarging, and connecting patches provided more moderate but comparatively consistent benefits across specialisation classes. Consequently, no single restoration strategy consistently maximized persistence benefits across the full trait space: strategies that strongly benefited specialists tended to underperform for generalists, and vice versa.

We also evaluated an alternative formulation in the Supplementary Information where restoration effort was standardized by the total area restored to maximum habitat quality, measured as the number of cells restored to *q_i_ = 1*, rather than effective habitat increase. Under this formulation, adding patches, enlarging patches, connecting patches, and improving habitat quality became more similar in efficiency, particularly for specialist species (S2 Fig. S2&S3). This suggests that specialists primarily benefited from the creation of high-quality habitat cells regardless of whether these arose through patch creation, expansion, corridors, or quality enhancement, whereas generalists remained more responsive to diffuse matrix improvement. Broad trait-dependent patterns were also qualitatively reflected in a complementary stochastic patch occupancy analysis (S3 Fig. S1).

## Discussion

Our work predicts that restoration outcomes depend on how interventions restructure habitat within landscapes. Under standardised restoration effort, different landscape restoration strategies produced substantially different gains in predicted persistence of species. These findings extend previous work showing that spatial prioritisation can determine where restoration delivers the greatest benefits across regions and landscapes (Strassburg *et al*. 2020) by demonstrating that, within a given landscape, the way restoration is implemented can itself strongly shape ecological outcomes. Rather than treating restoration as a uniform increase in habitat, our findings therefore suggest that different strategies should be understood as distinct ways of reshaping the effective landscape experienced by species, with their benefits depending on the traits of the species expected to respond.

The reason is that, in a metapopulation context, persistence depends not simply on the total amount of habitat available, but on how habitat quality and spatial arrangement combine to shape colonization and extinction across the landscape (Hanski 1999, 1998; Moilanen & Hanski 2001). The strategies considered here (Fig. 1A) therefore act on different ecological processes. Improving habitat quality reinforces existing patches; adding, enlarging, and connecting patches increase the area of suitable habitat in different spatial configurations; and matrix improvement changes the conditions through which individuals move or persist between habitat patches. Although these strategies may produce equivalent increases in effective habitat area, they do not create equivalent landscapes. Instead, they redistribute habitat quality and connectivity across the landscape network, altering where persistence potential is concentrated and which species are able to benefit from restoration.

This mechanistic distinction helps explain why persistence gains varied so strongly among restoration strategies (Fig. 3). Improving existing habitat quality was often among the most efficient strategy, because it concentrated restoration effort in parts of the landscape that already contributed to the habitat network. In the metapopulation capacity framework, persistence potential emerges from the network of pairwise connections among habitat cells, meaning that locally connected, high-quality habitat can have disproportionate influence (Hanski & Ovaskainen 2000; Moilanen & Hanski 2001). Quality improvement therefore does more than increase local suitability; it can strengthen many colonisation pathways simultaneously, especially within dense clusters of suitable habitat (Fletcher *et al*. 2013). Ecologically, this suggests that restoration may sometimes produce greater persistence gains by reinforcing strong habitat cores than by diffusely increasing suitability across the wider landscape. This interpretation aligns with work showing that the contribution of habitat to persistence depends on both its quality and its position within the wider landscape network (Mortelliti *et al*. 2010; Thomas *et al*. 2001; Wilson *et al*. 2023). However, this pattern should not be interpreted as a general ranking of restoration strategies as the apparent efficiency of different interventions depends strongly on which species are expected to respond.

Dispersal ability determined whether species could respond to restoration beyond their immediate neighbourhood, while specialisation determined which parts of the restored landscape were effectively usable. This creates a direct parallel with insights from investigations on habitat loss and fragmentation: just as species differ in their sensitivity to habitat loss and isolation because they perceive, use, and move through landscapes differently, they can also be expected to differ in their responses to restoration strategies that modified those same landscape properties (Ewers & Didham 2006; Henle *et al*. 2004; Suárez-Castro *et al*. 2026; Watts & Hughes 2024). Poor dispersers benefited little from most interventions because they remained constrained by spatial isolation, whereas more mobile species were better able to respond to changes in the wider habitat network (Fig. 4), consistent with work showing that dispersal limitation and effective isolation can restrict responses to habitat availability (Ricketts 2001; With & King 1999). Specialisation generated a second, distinct axis of variation: specialist species benefited disproportionately from improving habitat quality because their effective landscapes were restricted to high-quality habitat, whereas generalists derived greater benefit from matrix improvement because they could exploit more diffuse increases in usable area (Fig. 5). This supports a view of the matrix not simply as hostile non-habitat, but as a heterogeneous landscape component whose value depends on species’ habitat requirements and movement capacities (Driscoll *et al*. 2013; Fletcher *et al*. 2024; Kupfer *et al*. 2006; Ricketts 2001). Thus, the strong average performance of quality improvement partly reflects its benefits for specialists, while the low average performance of matrix improvement masks its potential value for generalists. These results reflect our representation of matrix improvement as increasing the availability of low-quality habitat; matrix restoration acting through other mechanisms, such as reducing mortality during dispersal, could generate different outcomes, including greater benefits for specialist species (Fletcher *et al*. 2026).

Because real landscapes support communities rather than single species, these trait-dependent responses imply that restoration strategies may reshape community composition through interspecific variation in species persistence. Habitat loss and fragmentation are widely understood to act as extinction filters (Betts *et al*. 2019), disproportionately excluding species whose traits make them sensitive to isolation, habitat degradation, or matrix hostility (Ewers & Didham 2006; Henle *et al*. 2004; Keddy 1992). Restoration might therefore be expected to reverse these filters, but our results suggest that different strategies are likely to relax different ecological constraints and therefore favour different trait groups during recolonisation and recovery, leading to ‘recovery filters’. Those recovery filters also point to a broader trade-off between concentrating restoration effort into a smaller amount of high-quality habitat and distributing it across a larger or more heterogeneous area. This is conceptually related to the area–heterogeneity trade-off, which proposes that increasing habitat heterogeneity may accommodate a broader range of ecological requirements while reducing the effective amount of suitable habitat available to any given species (Allouche *et al*. 2012; Ben-Hur & Kadmon 2020). Although habitat specialisation in the present model represents the degree of habitat quality required rather than specialisation across distinct habitat types, a similar tension emerges: concentrating restoration effort into high-quality habitat may particularly benefit species with stringent quality requirements, whereas more spatially diffuse interventions may generate a wider range of usable conditions but fail to create the extensive high-quality habitat cores required by such species. Restoration strategies may therefore differ not only in their average persistence benefits, but also in the trait filters they relax and the communities they are likely to favour. Biodiversity recovery at scale may require asking not only whether restoration increases persistence, but whether it addresses the particular filters that caused species to decline in the first place.

Our findings have direct implications for restoration planning under limited resources. First, they suggest efficiency might differ substantially among alternative spatial implementations of restoration (Fig. 3), reinforcing the idea that restoration at landscape scales is not a single action but a suite of mechanistically distinct interventions (Brudvig 2011; von Holle et al. 2020; Perring et al. 2015). In practice, selecting among such strategies is further complicated by the fact that restoration costs are rarely proportional to restored area or habitat gain, but instead vary strongly across intervention types and landscapes due to land values, feasibility constraints, and socio-economic context (Murdoch et al. 2007; Naidoo et al. 2006). This heterogeneity makes cost difficult to approximate in general simulation frameworks and implies that identifying optimal strategies will often require system-specific economic and logistical information. Second, our results highlight that restoration outcomes depend strongly on the traits of the species being targeted (Fig. 4–6), suggesting that strategy selection should explicitly consider traits of focal species such as dispersal ability and habitat specialization. While we focus on dispersal and specialisation as two key axes shaping landscape use, additional factors – such as demographic rates, behavioural movement responses, edge sensitivity, or competition – may further mediate restoration effectiveness and could shift the relative performance of alternative interventions (Ewers & Didham 2006; Henle et al. 2004). Third, pronounced specialist-generalist divergence indicates that restoration programs might not always be able to simultaneously maximize persistence gains across diverse communities using a single dominant intervention (Fig. 6). This tension emphasizes that restoration is inherently objective dependent, requiring explicit decisions about whether to prioritize for example specialist conservation, maximize overall persistence potential, or balance outcomes across functional groups.

While our framework provides general insights into how restoration efficiency may depend on spatial implementation and species traits, several limitations warrant consideration. First, we quantify restoration outcomes solely in terms of persistence potential, using metapopulation capacity as a mechanistic proxy (Hanski & Ovaskainen 2000). Persistence is a central conservation objective, but real restoration programmes often aim to deliver additional outcomes such as ecosystem functioning, resilience, carbon storage, or the recovery of ecological interactions (Bastin *et al*. 2019; Strassburg *et al*. 2020). Strategies that maximise persistence may therefore not necessarily maximise other restoration goals, highlighting the importance of aligning restoration design with explicit objectives. Second, the metric we focus on represents a long-term expectation of persistence potential and does not capture the transient dynamics through which restoration benefits are delivered (Hanski & Ovaskainen 2000). In practice, factors such as time lags in habitat recovery, delayed colonisation, extinction debt, and the sequence and timing of interventions may strongly influence both the magnitude and timing of observed restoration outcomes (Watts *et al*. 2020). Third, although we evaluate outcomes across trait space, we do not explicitly model multispecies dynamics. Species interactions such as competition, facilitation, or trophic dependencies could modify restoration responses in ways not captured by our single-species persistence metric (Leibold *et al*. 2004). Fourth, our simulated landscapes and interventions necessarily simplify real restoration contexts. In practice, restoration costs and feasibility vary spatially (Murdoch et al. 2007; Naidoo et al. 2006), habitat quality is multidimensional and dynamic (Allouche et al. 2012), and restored habitat may not achieve intended conditions (Suding 2011). Future work integrating empirical landscapes, spatial cost surfaces, and habitat recovery trajectories will be needed to assess how the trade-offs identified here translate into applied restoration planning.

Despite these simplifications, our results highlight a general principle: restoration strategies should be treated as mechanistically distinct ways of reshaping the effective landscape experienced by species. Equivalent gains in nominal habitat amount or quality can produce very different persistence outcomes depending on where they occur, how they alter connectivity, and which species are expected to respond. By linking restoration interventions to metapopulation theory and trait-dependent landscape perception, our framework generates testable predictions about when different restoration strategies should be most effective. Empirical studies that compare alternative restoration actions across landscapes, quantify changes in habitat quality and connectivity, and track responses of species differing in dispersal ability and habitat specialisation would provide a critical next step for evaluating these predictions. More broadly, our results suggest that successful landscape-scale restoration will require moving beyond area-based targets alone toward spatially explicit, trait-informed planning that asks not only how much habitat is restored, but what kind of habitat, where, and for whom.

## Acknowledgments

Robin Bauknecht acknowledges funding from the Gates Cambridge Trust and the National Environmental Research Council [grant number UKRI1346]. Robert Fletcher was supported by the Arcadia Foundation and the Endangered Landscapes and Seascapes Program.

