## Supplementary Information for "Landscape restoration strategies differ in their predicted benefits for species persistence"

**Authors and Affiliations:**

^1^ Department of Zoology, Conservation Research Institute, University of Cambridge, Cambridge, UK

**Table of Contents:**

**S1: Modelling Details**

**S1 Text S1:** Methodological details of restoration strategy implementation (pgs. 2-4)

**S1 Figure S1:** Simulating habitat landscapes (pg. 5)

**S1 Figure S2:** Visualizing landscape and habitat heterogeneity (pg. 6)

**S1 Figure S3:** Spatial implementation of specialisation function (pg. 7)

**S1 Figure S4:** Sensitivity to landscape heterogeneity (pg. 8)

**S1 Figure S5:** Sensitivity to habitat heterogeneity (pg. 9)

**S1 Figure S6:** Sensitivity to baseline habitat amount (pg. 10)

**S1 Figure S7:** Sensitivity to restoration effort (pg. 11)

**S1 Figure S8:** Frequency, magnitude, and influence of habitat-quality saturation on restoration outcomes (pg.12-13)

**S2: Alternative restoration-effort standardisation**

**S2 Text S1:** Methodological details of restoration strategy implementation under total-area standardisation (pgs. 14-15)

**S2 Figure S1:** Visualization of restoration under total-area standardisation (pg. 16)

**S2 Figure S2:** Average effects of landscape restoration strategies under total-area standardisation (pg. 17)

**S2 Figure S3:** Trait effects on restoration efficiency under total-area standardisation (pg. 18)

**S3: Stochastic Patch Occupancy Models (SPOMs)**

**S3 Text S1:** Dynamic simulation using SPOMs (pgs. 19-20)

**S3 Figure S1:** SPOM non-extinction probability across trait space (pgs. 21-22)

**S1: Modelling Details**

**S1 Text S1:** Methodological details of restoration strategy implementation

All restoration interventions operate on a spatial landscape object defined on a regular grid with (i) a binary habitat mask, (ii) realised habitat quality values $q_{i}$, and (iii) a latent spatial field describing habitat quality potential. Interventions modify habitat configuration and/or quality under a common restoration effort, defined as the increase in total effective habitat, $\Delta\sum_{i} q_{i}$ .

Across interventions, newly created cells receive quality values derived from the latent quality potential. Specifically, potential values at selected locations are (i) mapped monotonically into the bounded interval $\left[ q_{min}, q_{max} \right]$using rank-based scaling, and (ii) adjusted using a constrained projection algorithm to ensure that (a) all values remain within bounds and (b) their sum matches the specified effective habitat area increase. This procedure preserves spatial structure in the latent field while ensuring that the specified restoration effort is met exactly.

Below, we describe the algorithmic implementation of each intervention.

*Adding habitat patches*

The addition of new habitat patches is implemented by iteratively placing discrete patch geometries into the matrix. Patch shapes are drawn from an empirical library constructed from habitat patches observed in replicate landscapes of the same heterogeneity class. Shapes may be rotated and reflected to increase diversity while preserving realistic geometry.

At each iteration, the algorithm:

1. Estimates the typical effective habitat area increase associated with adding one cell, based on the mean quality of existing habitat.
2. Selects a candidate patch whose size is compatible with the remaining restoration effort.
3. Attempts to place the patch at a random location subject to three constraints: (i) all cells lie within the landscape, (ii) no overlap with existing habitat, and (iii) no direct 4-neighbor adjacency with existing habitat, enforced via a buffer.
4. Allocates quality values to the patch cells based on their latent potential, such that the resulting effective habitat area increase matches the amount allocated to that patch.
5. Updates the landscape and remaining restoration effort.

This process is repeated until no feasible placement is possible or the remaining restoration effort falls below the minimum required to create an additional habitat cell. Any residual effort can optionally be reallocated to improving existing habitat quality.

*Enlarging habitat patches*

Patch enlargement expands existing habitat outward into adjacent matrix cells. The algorithm first identifies all habitat patches as connected components (4-neighbour connectivity). It then performs a breadth-first expansion from all habitat cells simultaneously, assigning each matrix cell (i) a distance from the nearest habitat and (ii) an owning patch. Candidate cells are grouped by ring and by patch. Expansion proceeds ring-by-ring, and within each ring, cells are added in repeated sweeps such that each patch contributes at most one cell per sweep. This ensures balanced growth across patches.

For each batch of added cells:

- The number of cells is constrained by the remaining restoration effort.
- If necessary, only the highest-potential cells are retained.
- Quality values are assigned based on latent potential and scaled so that the resulting increase in effective habitat area matches the allocated amount.

Any remaining restoration effort can subsequently be allocated to uniform habitat quality improvement.

*Improving habitat quality*

Habitat quality improvement increases $q_{i}$ for existing habitat cells without altering spatial configuration. This is implemented using a constrained “water-filling” procedure:

1. The algorithm computes the available headroom for each cell, defined as the difference between its current value and the maximum $q_{max}$.
2. Restoration effort is distributed uniformly across all non-saturated habitat cells.
3. Cells reaching $q_{max}$ are removed from further updates, and remaining restoration effort is redistributed among the remaining cells.

This process continues until either the specified restoration effort is achieved or all habitat cells reach the maximum quality. If the latter occurs and effort remains, the excess can optionally be redirected to habitat expansion.

*Improving matrix quality*

Matrix improvement increases the quality of all non-habitat cells uniformly. The specified restoration effort is distributed evenly across all matrix cells, such that each cell receives an identical increment,

$$\Delta q=\frac{overall effective habitat increase}{N_{matrix}}$$

where $N_{matrix}$ is the number of matrix cells. This intervention effectively transforms the matrix into low-quality, spatially homogeneous habitat.

*Connecting habitat patches*

Disconnected habitat components are connected through directed placement of new habitat. The algorithm proceeds iteratively:

1. Habitat patches are identified and their boundary cells extracted.
2. A multi-source breadth-first search (BFS) is initiated simultaneously from all patch boundaries, propagating through matrix cells.
3. When wavefronts originating from different patches meet, the shortest connecting path between those patches is identified.
4. The path is reconstructed and optionally widened by applying a local buffer.
5. A subset of corridor cells is selected based on the remaining restoration effort, and quality values are assigned according to latent potential.

This process is repeated until all patches are connected or the specified restoration effort is achieved. If connectivity is achieved before the specified restoration effort is achieved, remaining resources can be used to widen the corridor network by expanding outward from the connected structure. Any residual restoration effort can optionally be reallocated to improving habitat quality.

All algorithms are implemented in R and are fully reproducible. The complete implementation, including all helper functions and visualisation workflows, is provided in the accompanying code repository.

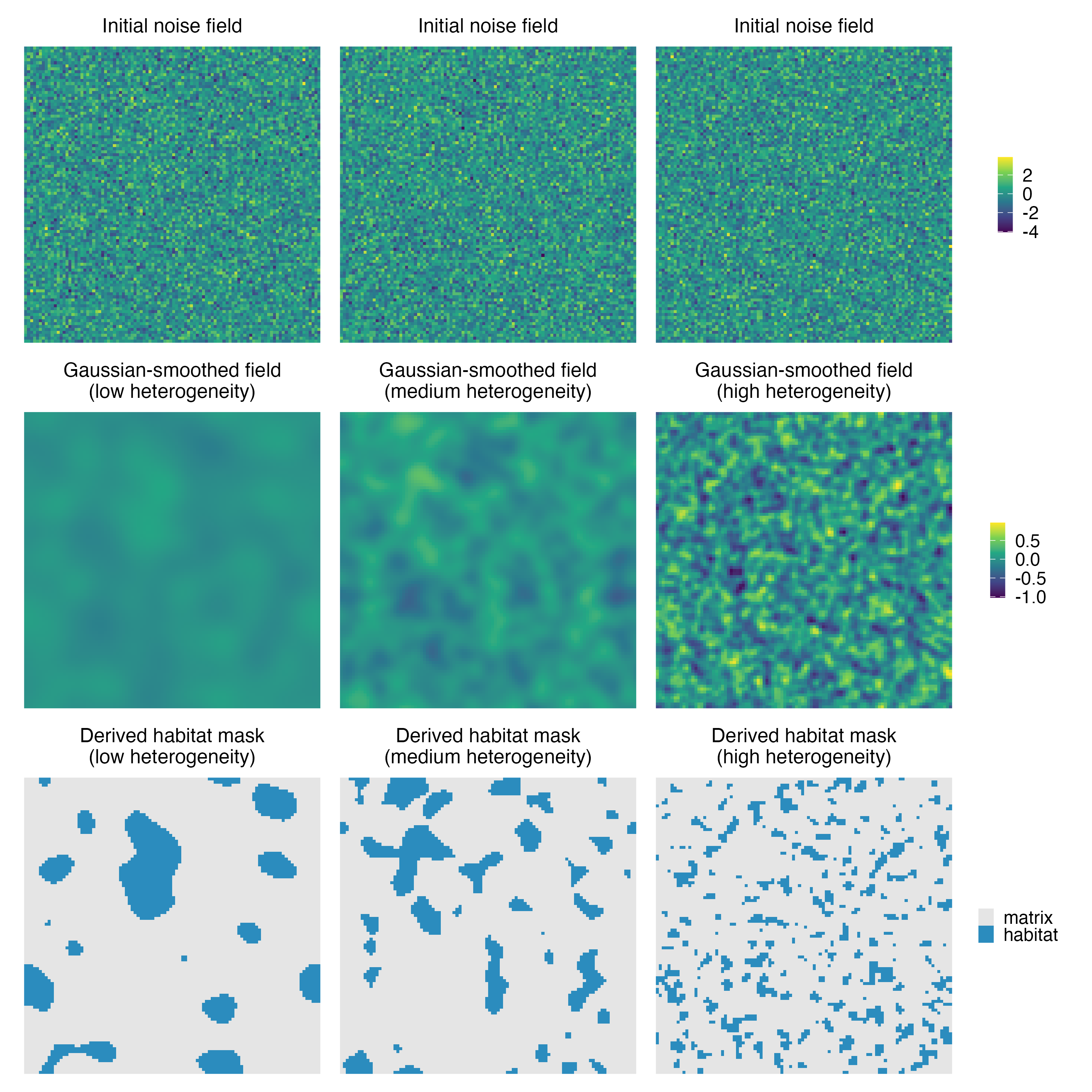

**S1 Figure S1 | Simulating habitat landscapes**

Top row: independent Gaussian noise fields. Middle row: the same fields after Gaussian smoothing with kernel width ($\sigma$), producing low ($\sigma_{L}=5$), medium ($\sigma_{L}=2.5$), and high ($\sigma_{L}=1$) landscape heterogeneity from left to right. Bottom row: binary habitat masks obtained by thresholding the smoothed fields to a fixed habitat amount (here 1500 cells; 15% of landscape); habitat patches emerge as contiguous clusters (blue), with matrix shown in grey. A second, independent smoothed field (not shown here, see compiled code for visualization) was generated for each landscape and treated as the latent habitat‑quality potential, which was mapped onto the habitat masks to assign within‑patch quality.

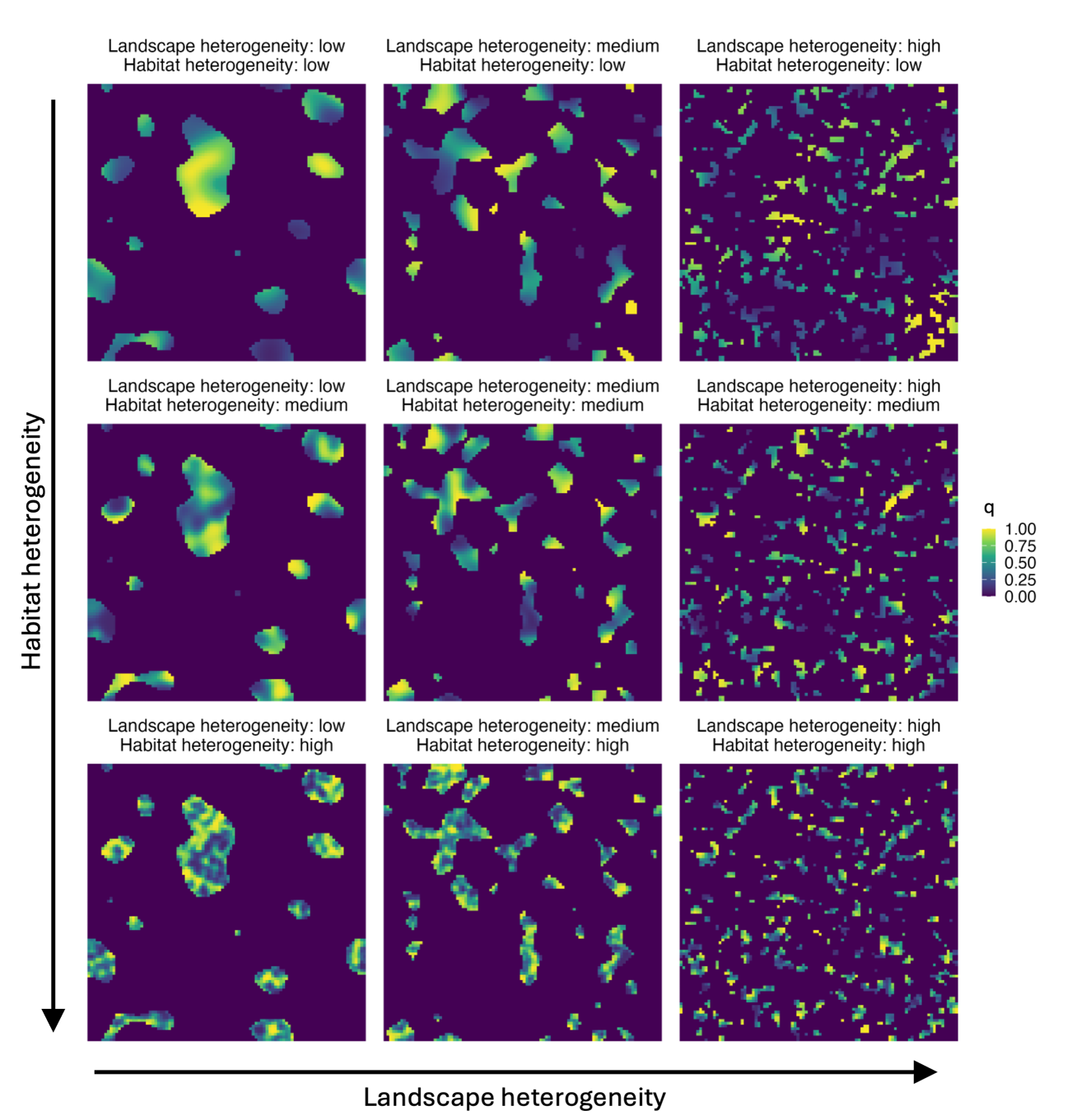

**S1 Figure S2 | Visualizing landscape and habitat heterogeneity**

Example landscapes illustrating independent variation in landscape heterogeneity (columns: low, medium, high; $\sigma_{L}=5, 2.5, 1$) and within‑patch habitat heterogeneity (rows: low, medium, high; $\sigma_{q}=5, 2.5, 1$). Habitat amount is fixed (here 1500 cells; 15% of the landscape), so differences among panels reflect spatial arrangement of habitat and within‑patch quality structure. Colors show habitat quality $q$.

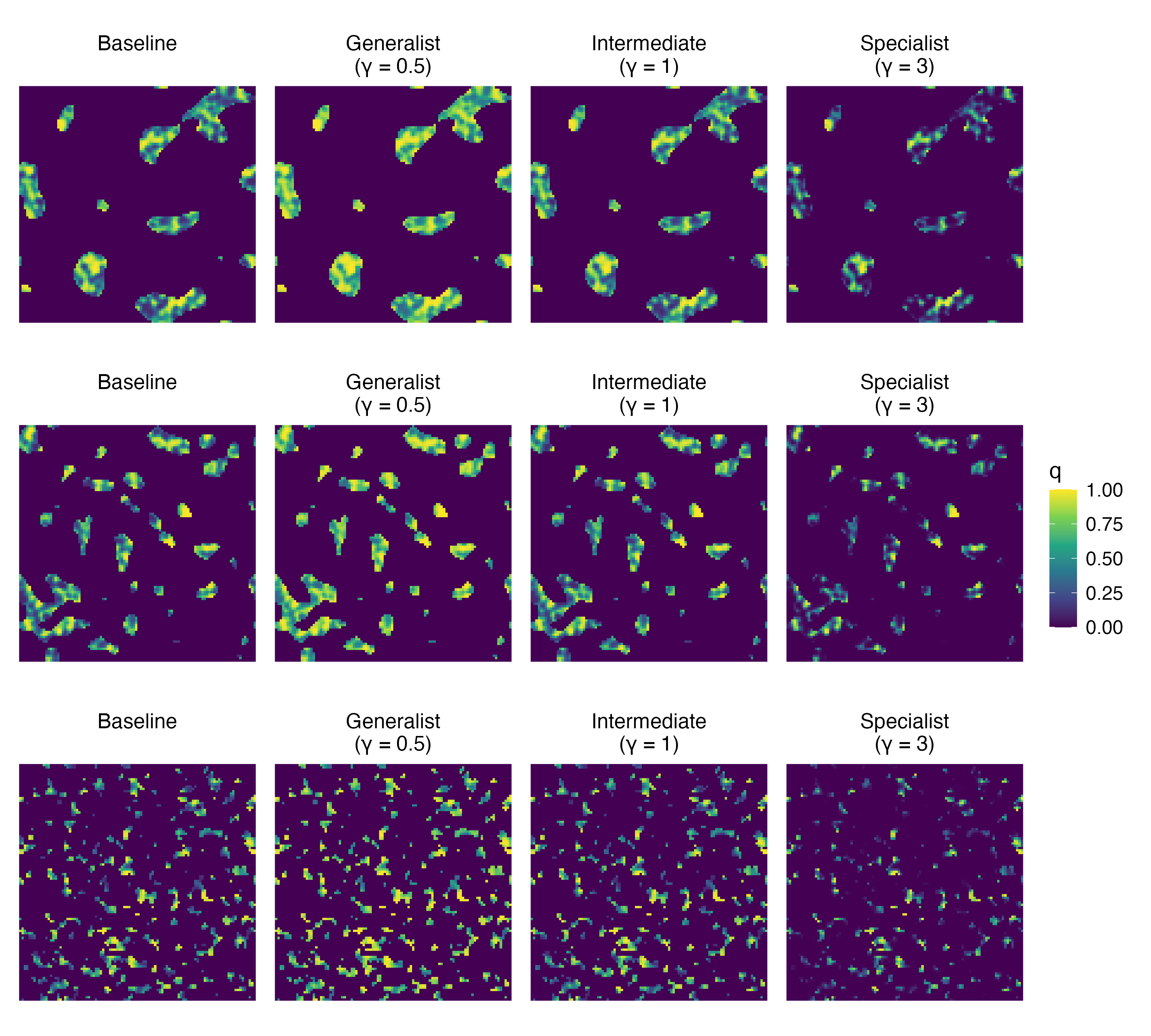

**S1 Figure S3 | Spatial implementation of specialisation function**

Example landscapes (rows: low, medium, high landscape heterogeneity, $\sigma_{L}=5, 2.5, 1$, respectively) shown at baseline habitat quality $q$ (left column) and after applying the specialisation transformation $q^{\gamma}$ for a generalist ($\gamma=0.5$), intermediate ($\gamma=1$), and specialist ($\gamma=3$) species. Increasing $\gamma$ accentuates high‑quality habitat and down‑weights low‑quality habitat, while preserving the spatial configuration of habitat patches.

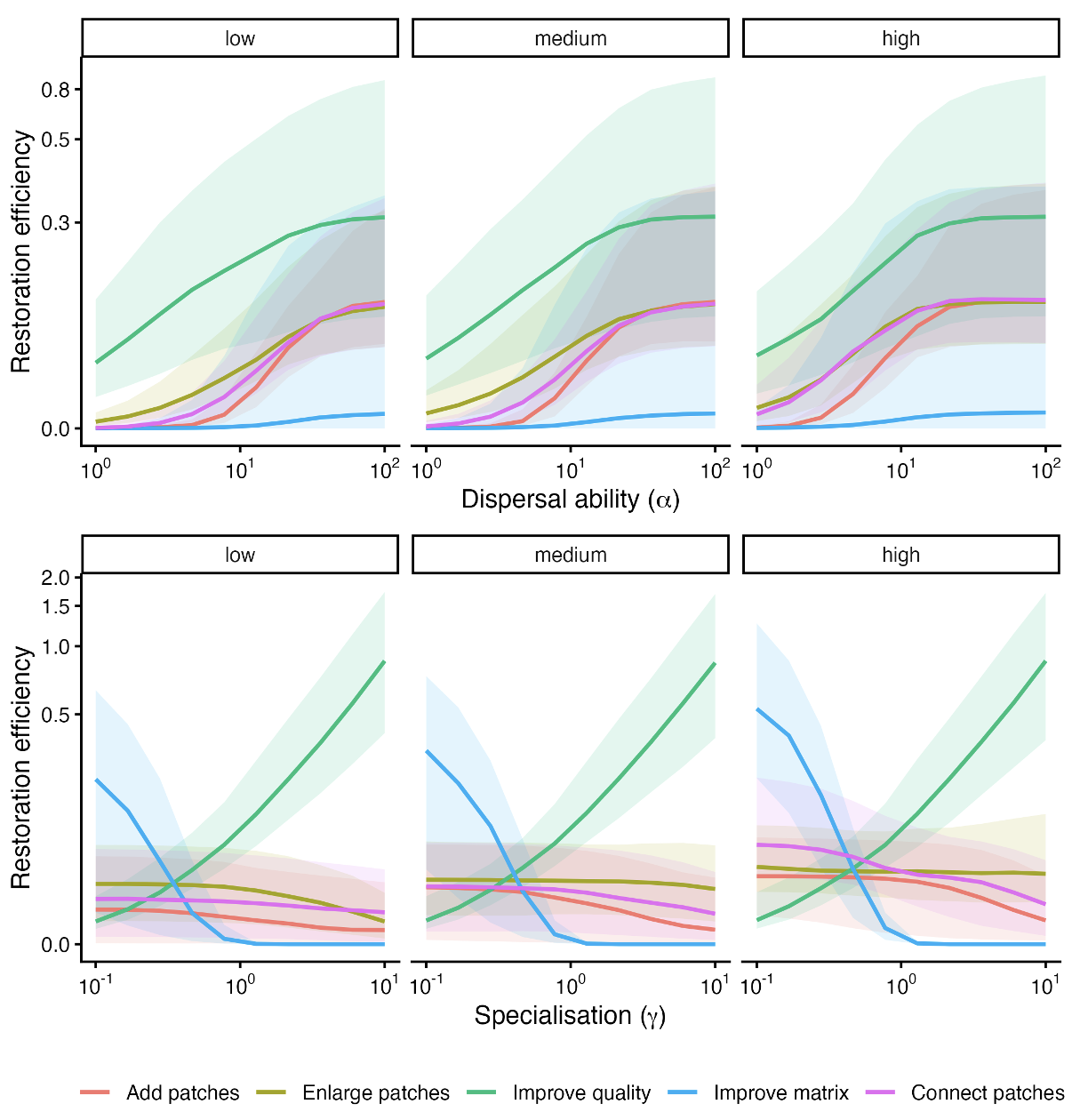

**S1 Figure S4 | Sensitivity to landscape heterogeneity**

**(A)** Restoration efficiency–α and **(B)** restoration efficiency–γ relationships, shown separately for low, medium, and high landscape heterogeneity (low, medium, high landscape heterogeneity, $\sigma_{L}=5, 2.5, 1$, respectively). Lines show median restoration efficiency and ribbons the interquartile range. The qualitative patterns and relative ordering of interventions are consistent across heterogeneity levels, indicating that the trait–restoration efficiency relationships are robust to variation in patch configuration.

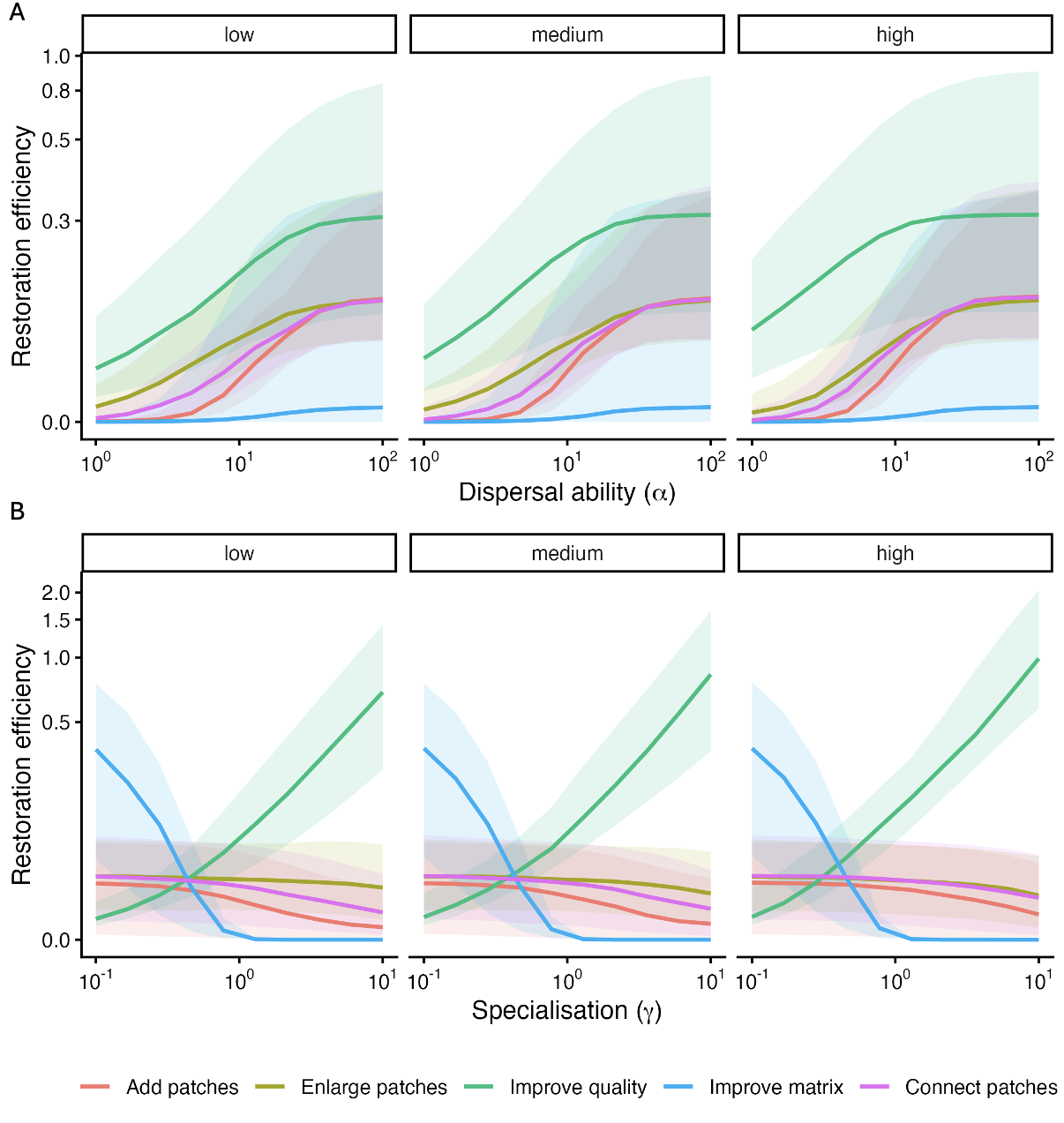

**S1 Figure S5 | Sensitivity to habitat heterogeneity**

**(A)** Restoration efficiency–α and **(B)** restoration efficiency–γ relationships, shown separately for low, medium, and high landscape heterogeneity (low, medium, high landscape heterogeneity, $\sigma_{q}=5, 2.5, 1$, respectively). Lines show median restoration efficiency and ribbons the interquartile range. The qualitative patterns and relative ordering of interventions are consistent across heterogeneity levels, indicating that the trait–restoration efficiency relationships are robust to variation in within-patch habitat configuration.

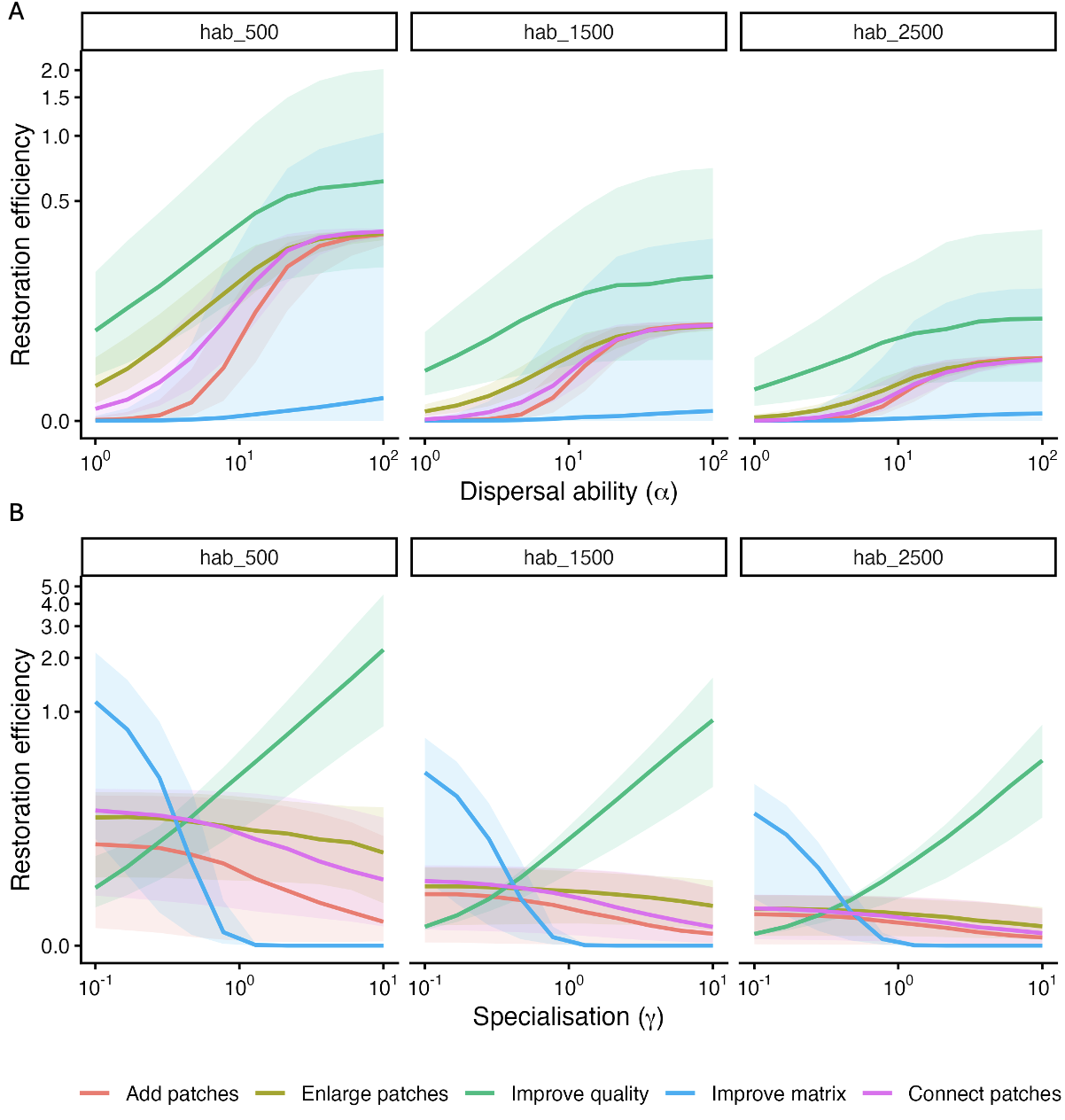

**S1 Figure S6 | Sensitivity to baseline habitat amount**

**(A)** Restoration efficiency–α and **(B)** restoration efficiency–γ relationships shown for landscapes with low (500 cells total habitat, “hab_500”), intermediate (1500 cells total habitat, “hab_1500”), and high (2500 cells total habitat, “hab_2500”) total habitat. Lines show medians and ribbons show interquartile ranges. Landscapes with less initial habitat tend to show larger proportional gains (higher restoration efficiency), consistent with lower baseline metapopulation capacity amplifying percentage improvements. However qualitatively the relationships are similar across landscapes with varying total habitat.

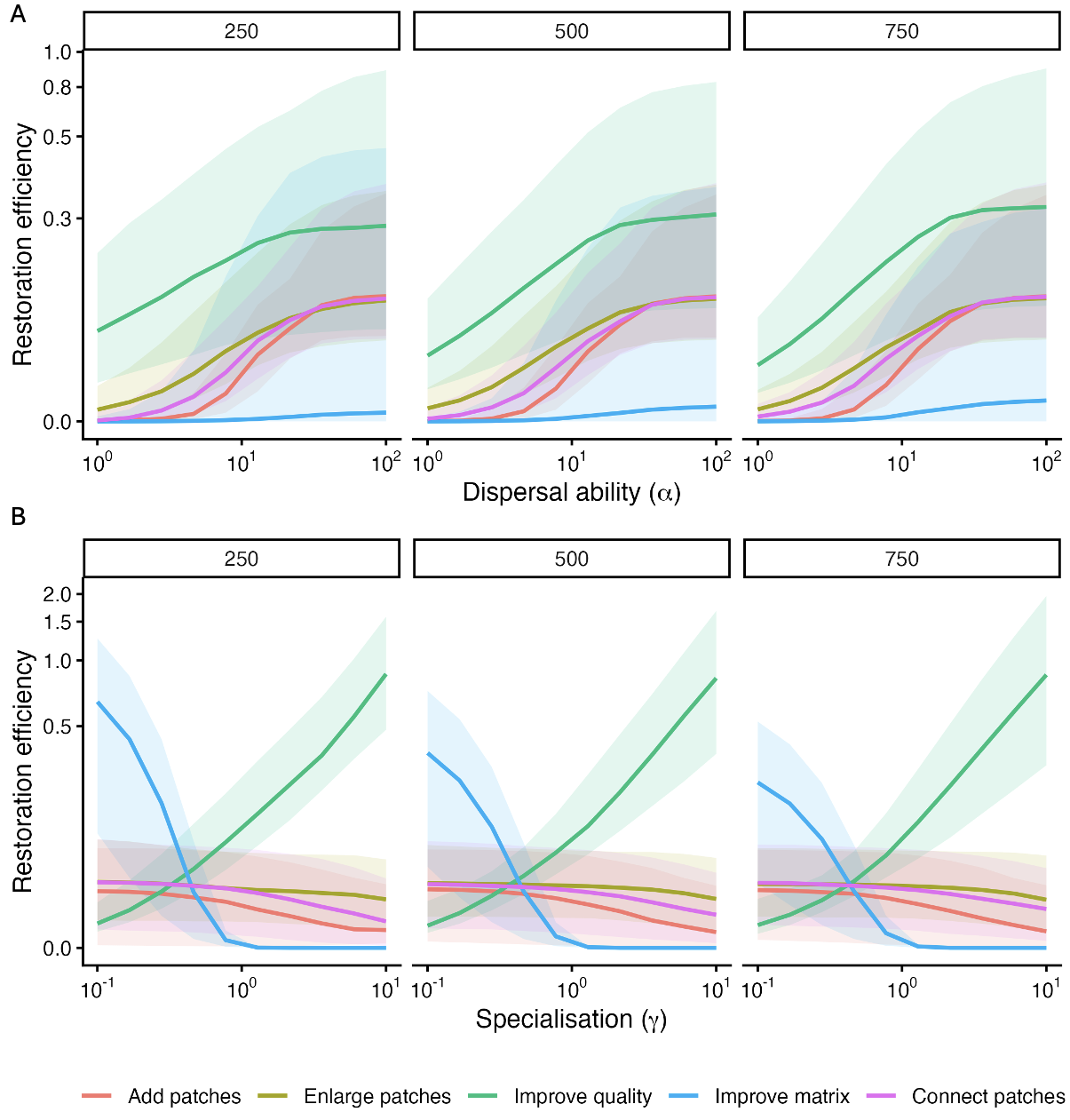

**S1 Figure S7 | Sensitivity to restoration effort**

**(A)** Restoration efficiency–α and **(B)** restoration efficiency–γ relationships shown for three levels of restoration effort ($\Delta\sum_{i} q_{i}=250, 500, 750$). Lines show medians and ribbons the interquartile range. Curves are highly similar across restoration-effort levels, supporting the use of restoration efficiency, $\%\Delta\lambda$ / effective habitat area restored) as a comparable efficiency metric.

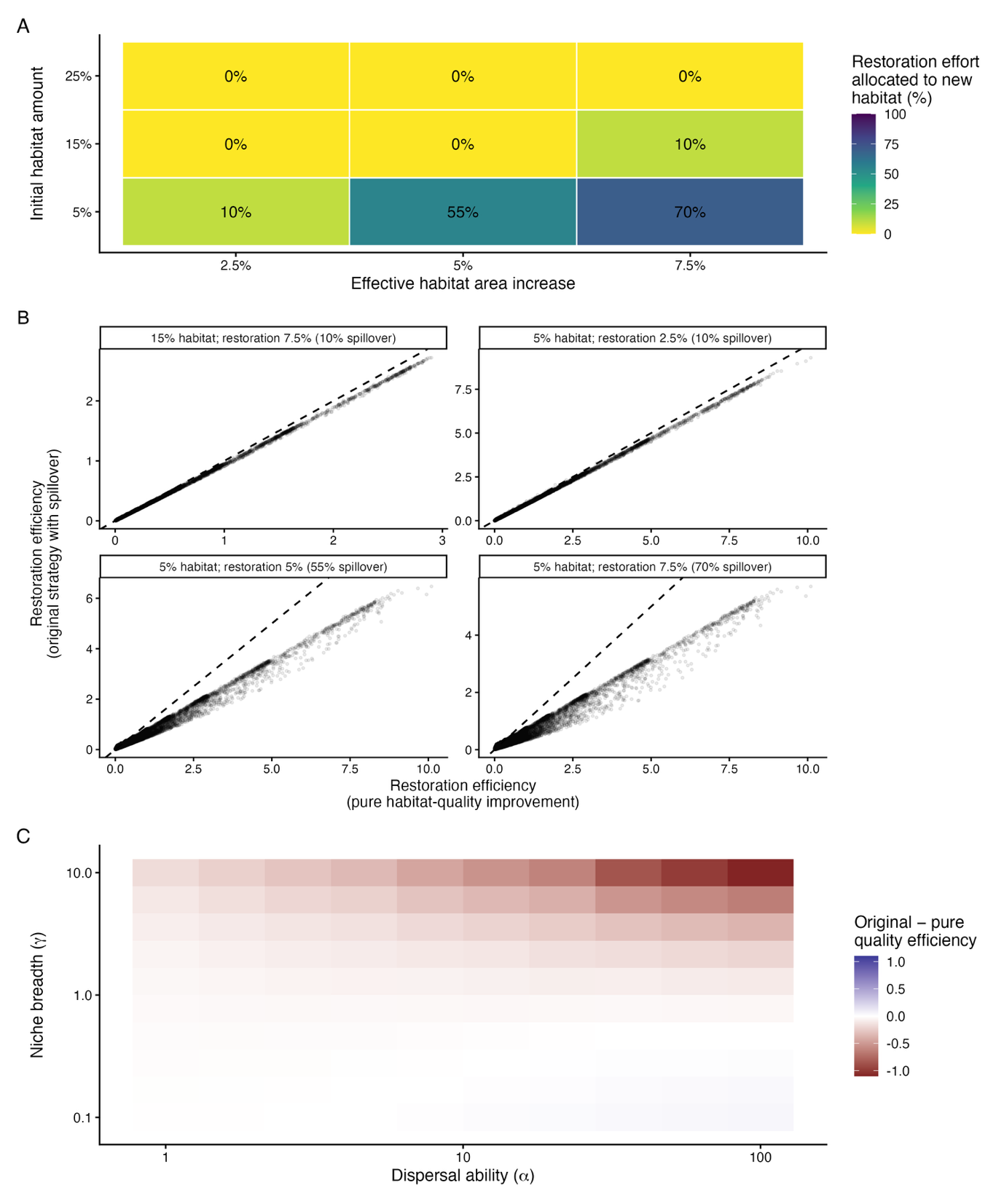

**Figure S8 | Frequency, magnitude, and influence of habitat-quality saturation on restoration outcomes
(A)** Proportion of the prescribed effective habitat area increase that was implemented as newly created habitat after all existing habitat cells had reached maximum quality under the habitat-quality improvement strategy, across combinations of initial habitat amount and prescribed effective habitat area increase. Spillover occurred when the prescribed increase exceeded that achievable through improvement of existing habitat alone. Allowing spillover therefore enabled all landscape–restoration combinations to retain the same prescribed increase in effective habitat area as the other restoration strategies. **(B)** Matched comparison of restoration efficiency immediately after all existing habitat reached maximum quality (“pure habitat-quality improvement”; x-axis) and after the remaining prescribed restoration effort had additionally been implemented through expansion into adjacent matrix (“original strategy with spillover”; y-axis). Each point represents the same landscape and trait combination, and the dashed line denotes equal restoration efficiency. The saturation-only comparator necessarily uses less total restoration effort than the original strategy and therefore does not represent an equal-effort alternative. Points below the 1:1 line indicate that restoration implemented after quality saturation had lower marginal returns than the preceding quality-improvement phase, thereby reducing average restoration efficiency across the full intervention. The results show that the high restoration efficiency of habitat-quality improvement in the main analysis was not generated by subsequent habitat expansion; rather, spillover generally diluted the efficiency achieved during the initial quality-improvement phase. **(C)** Median difference in restoration efficiency between the original implementation including spillover and the saturation-only habitat-quality improvement comparator across dispersal ability ($\alpha$) and specialisation ($\gamma$). Negative values indicate that continuing restoration through adjacent habitat creation after quality saturation reduced average restoration efficiency relative to stopping at saturation. This reduction was strongest for highly mobile specialist species, corresponding to the region of trait space in which habitat-quality improvement showed its strongest relative performance in the main analysis. Spillover therefore did not generate the strong specialist response to habitat-quality improvement and instead dampened it, supporting the robustness of the main conclusions to the spillover implementation.

**S2: Alternative Budget Formulation**

**S2 Text S1:** Methodological details of restoration strategy implementation under cell-based budget formulation

In the alternative formulation, restoration effort was standardised by the total area restored to maximum habitat quality, rather than by the increase in effective habitat area. One unit of restoration effort therefore corresponded to one cell restored to maximum habitat quality, and interventions were evaluated at total restored areas of 250, 500, and 750 cells.

Under this definition, the matrix improvement intervention was not included, because uniformly increasing matrix quality across all non-habitat cells does not correspond naturally to a discrete number of fully restored cells. The cell-based analysis therefore considered only four interventions: adding habitat patches, enlarging habitat patches, improving habitat quality, and connecting habitat patches.

Under this formulation, restored cells were generally assigned maximum habitat quality $q=1$ rather than receiving continuous quality values scaled to match a target expenditure. As a result, the role of the latent quality potential differed from that in the main analysis. Instead of determining the exact quality assigned to restored cells, latent potential was used primarily to prioritise which candidate cells were restored when more eligible cells were available than could be accommodated within the remaining number of cells to be restored.

Below, we describe the algorithmic implementation of each intervention under this alternative standardisation.

*Adding habitat patches*

The addition of new habitat patches is implemented by iteratively placing discrete patch geometries into the matrix. As in the main analysis, patch shapes are drawn from an empirical library constructed from habitat patches observed in replicate landscapes of the same heterogeneity class. Shapes may be rotated and reflected to increase diversity while preserving realistic geometry.

At each iteration, the algorithm:

1. Selects a candidate patch whose size is compatible with the remaining number of cells that can be restored.
2. Attempts to place the patch at a random location subject to three constraints: (i) all cells lie within the landscape, (ii) no overlap with existing habitat, and (iii) no direct contact (4-neighbour adjacency) with existing habitat, enforced via a buffer.
3. If placement succeeds, all cells in the patch are converted to habitat and assigned maximum quality ($q=1$).
4. The remaining number of cells to be restored is reduced by the number of cells added.

If no patch shape can be placed within the remaining number of cells, the algorithm falls back to selecting individual eligible cells. In this case, cells with the highest latent quality potential are prioritised and set to $q=1$.

*Enlarging habitat patches*

Patch enlargement expands existing habitat outward into adjacent matrix cells. The algorithm first identifies all habitat patches as connected components under 4-neighbour connectivity. It then performs a breadth-first expansion from all habitat cells simultaneously, assigning each matrix cell (i) a distance from the nearest habitat and (ii) an owning patch. Candidate cells are grouped by ring and by patch. Expansion proceeds ring-by-ring, and within each ring, cells are added in repeated sweeps such that each patch contributes at most one cell per sweep. This ensures balanced growth across patches. All selected cells are converted to habitat and assigned maximum quality ($q=1$). If more candidate cells are available within a sweep than the remaining number of cells to be restored, cells with the highest latent quality potential are prioritised.

Improving habitat quality

Habitat quality improvement increases the quality of existing habitat cells without altering spatial configuration. Under total-area standardisation, this intervention does not distribute incremental quality increases across all habitat cells. Instead, it selects a fixed number of existing habitat cells and upgrades them directly to maximum quality ($q=1$). Eligible cells are existing habitat cells with $q<1$. These are ranked first by their current quality, such that cells with lower existing quality are prioritised, and second by latent quality potential as a tie-breaker. The selected cells are then set to $q=1$ u until the specified number of cells has been restored or no eligible cells remain.

*Connecting habitat patches*

Disconnected habitat components are connected through the directed placement of new habitat cells.

The algorithm proceeds iteratively:

1. Habitat patches are identified and their boundary cells extracted.
2. A multi-source breadth-first search (BFS) is initiated simultaneously from all patch boundaries, propagating through matrix cells.
3. When wavefronts originating from different patches meet, the shortest connecting path between those patches is identified.
4. The path is reconstructed and optionally widened by applying a local buffer.
5. Newly selected corridor cells are converted to habitat and assigned maximum quality ($q=1$).

This process is repeated until all patches are connected or the specified number of restored cells is reached. If connectivity is achieved before the specified number of cells has been restored, remaining restoration effort can be used to widen the connected corridor network outward from the existing structure. During this widening step, if the number of candidate cells exceeds the remaining number of cells to be restored, cells with the highest latent quality potential are prioritised.

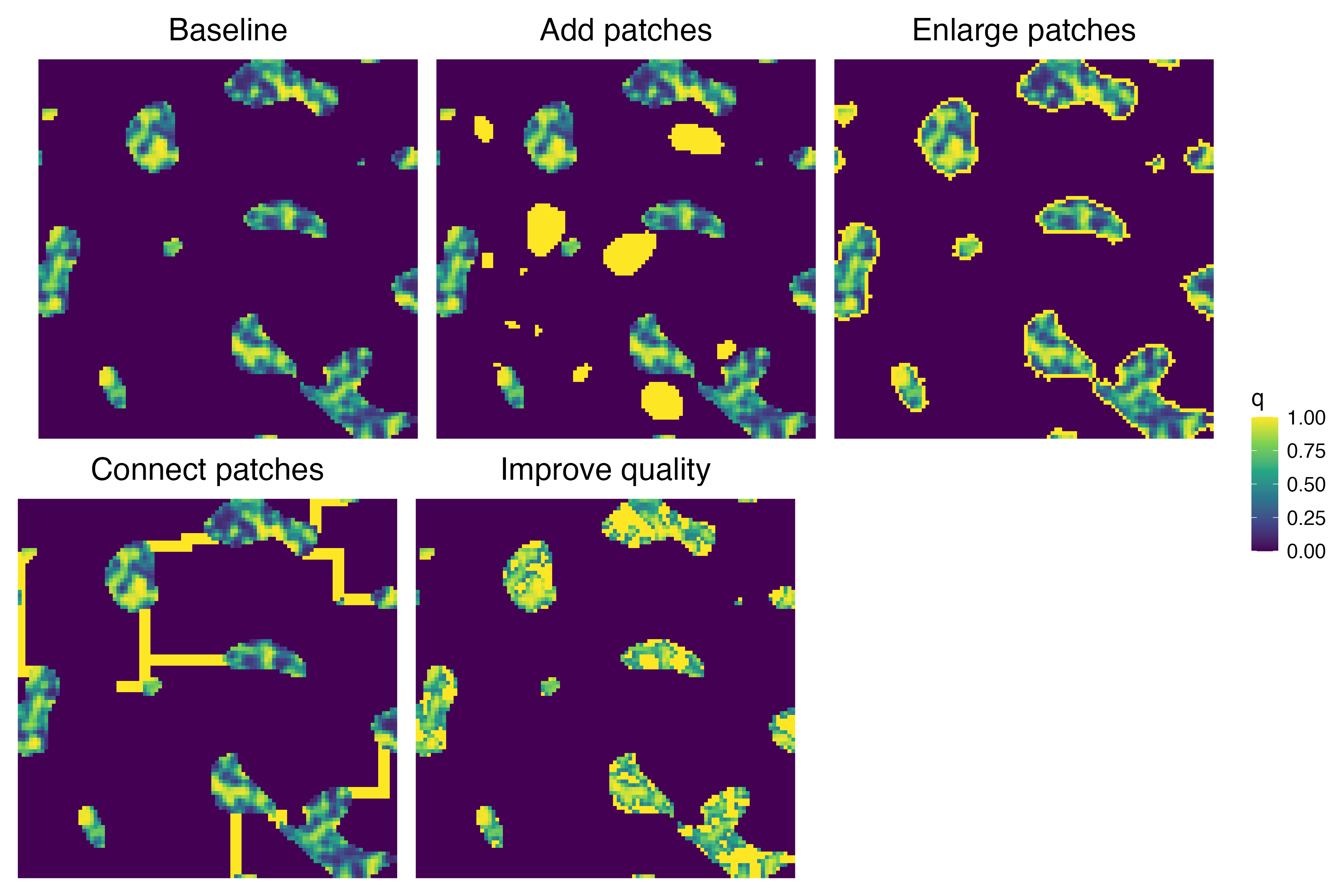

**S12 Figure S1 | Visualization of restoration under total-area standardisation**

Example landscape showing restoration outcomes under the alternative total-area standardisation. A baseline landscape is shown alongside four restoration strategies: adding patches, enlarging patches, connecting patches, and improving habitat quality. In this formulation, restoration effort is standardised by the number of cells restored to maximum habitat quality; here, 500 cells. Matrix improvement is omitted as described in S2 Text S1.

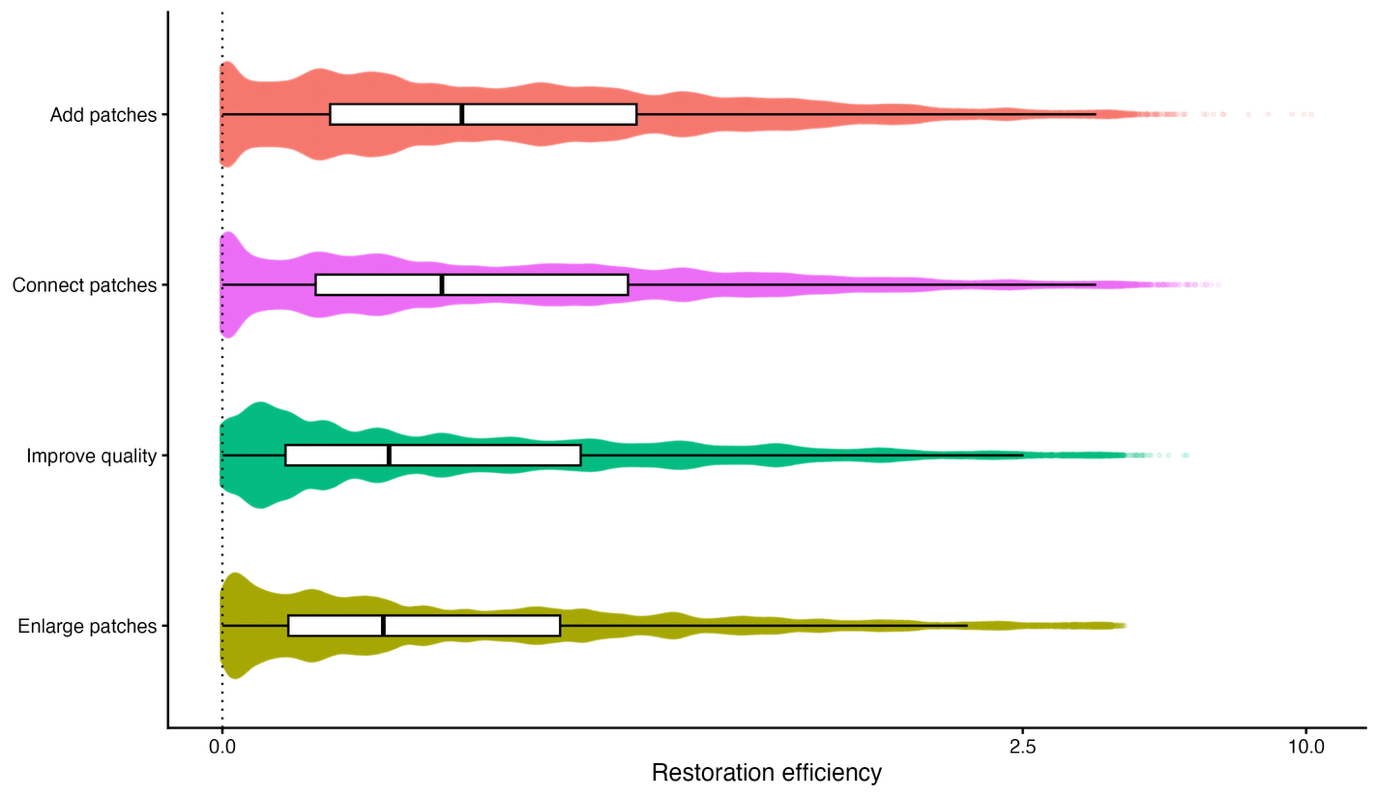

**S2 Figure S2 | Average effects of landscape restoration strategies under total-area standardisation**

Distribution of restoration efficiency under the alternative total-area standardisation. Restoration efficiency was calculated as the signed percentage change in metapopulation capacity per restored cell ($\%\Delta\lambda$ / restored cell). Scenarios span the full factorial set of simulated landscapes, landscape and habitat heterogeneity, replicate landscapes, restoration-effort levels, and the full dispersal-specialisation trait grid. Violin plots show kernel density; boxes indicate the interquartile range and the central line is the median. Under this formulation, adding patches, enlarging patches, connecting patches, and improving habitat quality showed broadly overlapping values of restoration efficiency, indicating that creating high-quality habitat cells can generate similar persistence benefits across different spatial arrangements when restored cells reach maximum quality. Matrix improvement is omitted because the alternative analysis focused on interventions that create or upgrade habitat cells to maximum quality.

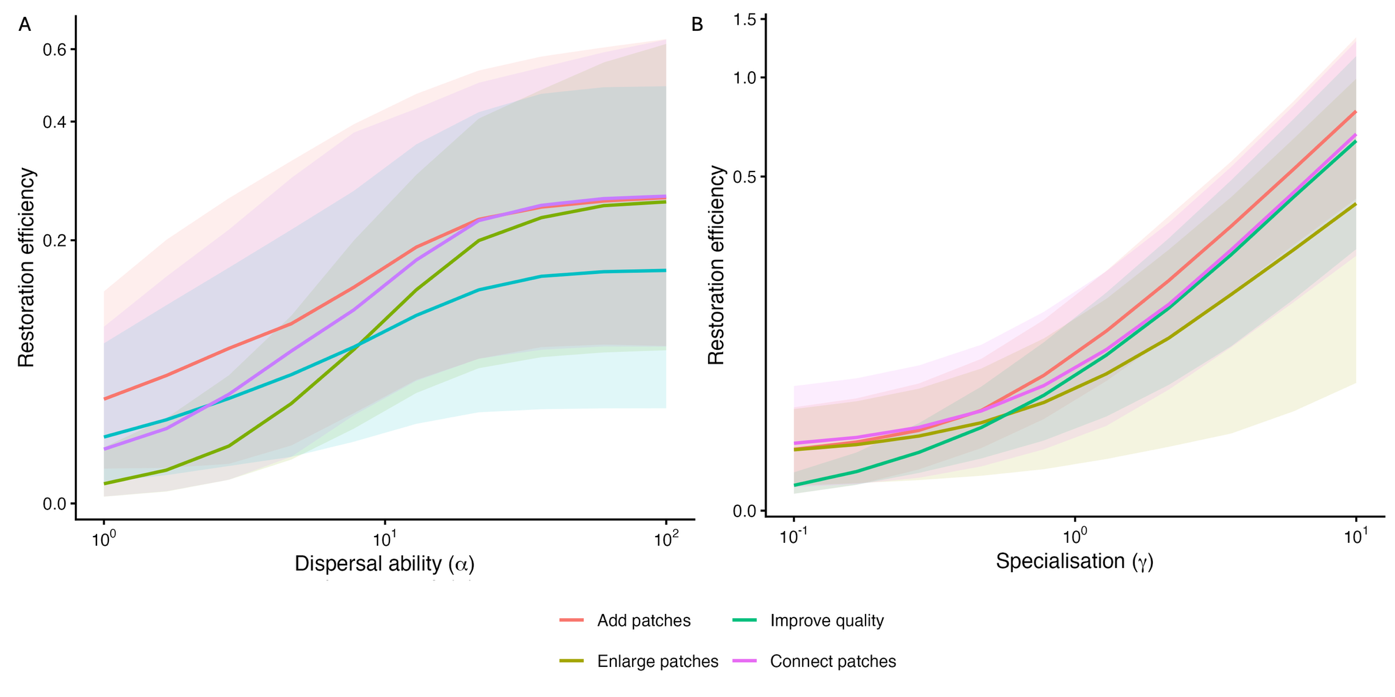

**S2 Figure S3 | Trait effects on restoration efficiency under total-area standardisation**

Restoration efficiency of restoration strategies under the alternative total-area standardisation. Restoration efficiency was calculated as the signed percentage change in metapopulation capacity per restored cell ($\%\Delta\lambda$ / restored cell). Panels show how restoration efficiency varied with dispersal ability $\alpha$ and specialisation $\gamma$, averaged across simulated landscapes, landscape and habitat heterogeneity, replicate landscapes, and restoration-effort levels. Lines indicate median restoration efficiency, and shaded ribbons indicate interquartile ranges. Under this alternative standardisation, adding patches, enlarging patches, connecting patches, and improving habitat quality showed broadly similar trait-response curves. This indicates that specialists benefited primarily from the creation of high-quality habitat cells, regardless of whether those cells were created through new patches, patch expansion, corridors, or quality enhancement.

**S3: Stochastic Patch Occupancy Models (SPOMs)**

**S3 Text S1:** Dynamic simulation using SPOMs

**Stochastic patch occupancy simulations on pixel-based landscapes**

To complement the metapopulation-capacity analysis, we evaluated restoration effects using a stochastic patch occupancy model (SPOM) implemented directly on the pixel-based landscapes (Hanski 1994, 1998). While metapopulation capacity provides a deterministic proxy for long-term persistence potential (Hanski & Ovaskainen 2000), the SPOM allowed us to assess whether restored landscapes produced similar trait-dependent differences in explicit stochastic dynamics through time.

*Landscape selection and restoration scenarios*

Rather than replicating the full factorial analysis, we used a single intermediate landscape scenario to test whether the main patterns were qualitatively reproduced under explicit stochastic dynamics. We selected a non-extreme configuration that allowed all restoration strategies to express meaningful effects while remaining broadly representative of the simulated landscape set. Specifically, the baseline landscape contained 1500 habitat cells (15% of the landscape), medium landscape heterogeneity ($\sigma_{L}=2.5$), high within-patch habitat-quality heterogeneity ($\sigma_{q}=1$), and replicate identity 1. From this baseline, we generated one restored landscape for each restoration strategy using the same intervention functions as in the main analysis and a restoration effort corresponding to an effective habitat area increase of 500 units under the primary standardisation. This resulted in six landscape states: baseline, add patches, enlarge patches, connect patches, improve habitat quality, and improve matrix quality.

*Trait parameterization*

Each landscape cell was treated as a discrete unit with a binary occupancy state (occupied or unoccupied), and cells with positive habitat quality in a given landscape state were considered available for occupancy. For each species trait combination, realised habitat quality $q_{i}$ were transformed into effective habitat quality as

$$q_{i}^{eff}=q_{i}^{\gamma}$$

where $\gamma$ controls specialisation. Low $\gamma$ values correspond to generalists (compressed quality differences), whereas high γ values correspond to specialists (greater weighting of high-quality cells). To maintain comparability with the metapopulation-capacity analysis while keeping simulations tractable, we evaluated three representative values each of dispersal ability ($\alpha$) and specialisation ($\gamma$), corresponding to low, intermediate, and high positions along the full trait gradients.

*Simulation dynamics*

Occupancy dynamics were simulated in discrete annual time steps. To ensure that differences among restoration strategies reflected changes in landscape configuration and quality rather than differences in initial population states, each baseline and restored landscape was initialized from the same pre-restoration occupancy pattern. At the start of each stochastic replicate, cells with positive effective habitat quality in the baseline landscape were independently initialized as occupied with probability$P_{init}=0.5$. The identical initial occupancy realization was then applied to the baseline and all five restored landscape states. Consequently, cells made newly occupiable through restoration were initially unoccupied and could contribute to persistence only after being reached through the colonisation process.

Each annual time step consisted of two sequential processes: local extinction followed by colonisation.

Extinction:

Occupied cells underwent local extinction with probability:

$$P_{ext,i}=1-exp(-\frac{e}{q_{i}^{eff}+\varepsilon})$$

where $e$ is a baseline extinction parameter and $\varepsilon$ is a small constant included for numerical stability. Under this formulation, extinction risk is highest in low-quality cells and declines as habitat quality increases.

Colonization:

Empty occupiable cells could be colonized by propagules originating from occupied cells. Dispersal followed an exponential kernel:

$$K\left( d_{ij} \right)=exp(-d_{ij}/\alpha)$$

The kernel was normalized to sum to one, so that $\alpha$ primarily controlled the spatial distribution of colonization pressure while its overall magnitude was governed by the colonization parameter $c$. This differs from the metapopulation-capacity analysis, where the kernel was not normalized and dispersal directly influenced total connectivity.

Colonization pressure at each cell i was calculated as:

$$S_{i}=\sum_{j} K\left( d_{ij} \right)q_{j}^{eff}O_{j}$$

where $O_{j}$ is the occupancy state of source cell $j$. For each dispersal ability–specialisation combination, colonisation pressure was rescaled relative to the corresponding baseline landscape:

$$S_{i}^{scaled}=\frac{S_{i}}{S_{ref}}$$

where $S_{ref}$ is the median colonization pressure in the baseline landscape under full occupancy for the corresponding $\alpha$- $\gamma$ combination. Because the absolute magnitude of $S_{i}$ can varies strongly across landscapes and trait combinations, unscaled values can lead to trivial dynamics (near-zero or near-certain colonization). Rescaling ensures that the colonization parameter operates on a comparable scale and maintains non-trivial dynamics across scenarios. Colonization probability was then given by:

$$S_{col,i}=1-exp(-cS_{i}^{scaled})$$

*Simulation design and response variable*

For each combination of dispersal ability, specialisation, colonisation parameter, and extinction-pressure parameter, we generated 100 stochastic replicates. Replicates were paired across the baseline and five restoration states: corresponding states shared the same initial occupancy realization and the same underlying cell- and time-specific random draws for extinction and colonisation. The colonization parameter $c$ was varied across 10 evenly spaced values between 0 and 1, and the extinction parameter $e$ was evaluated at three levels ($e=0.25, 0.5, 0.75$). Simulations were run for up to 25 annual time steps, with global extinction treated as an absorbing state. For each replicate, we recorded whether global extinction had occurred by years 5, 10, and 25. The primary response variable was the probability of non-extinction by time $T$:

$$1-P(extinct by T)$$

estimated as the proportion of 100 stochastic replicates in which at least one cell remained occupied at the specified time horizon.

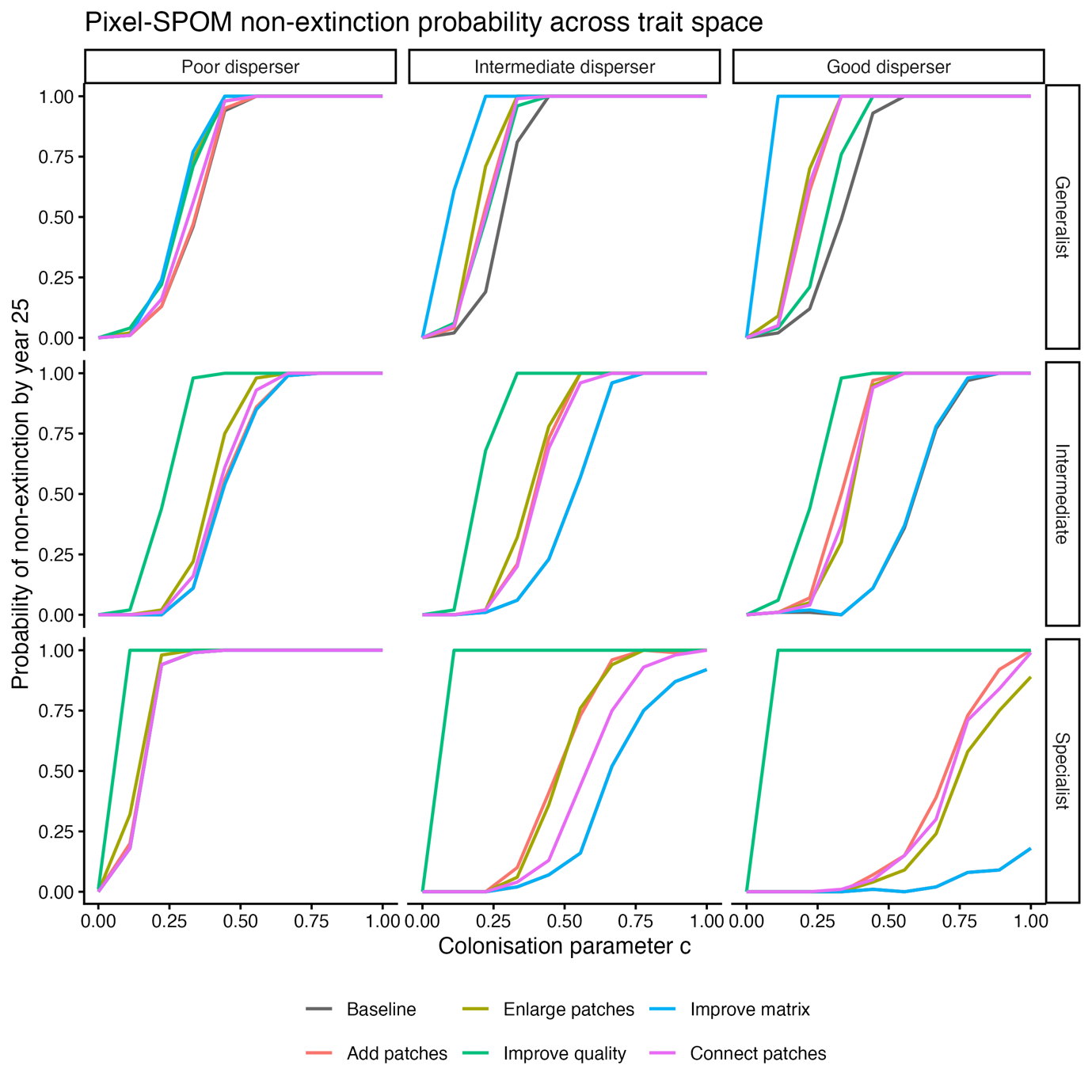

**S3 Figure S1 | SPOM non-extinction probability across trait space**

Stochastic patch occupancy model (SPOM) results for one focal restoration scenario. Curves show the estimated probability of non-extinction by year 25 as a function of the colonisation parameter $c$ for the baseline landscape and each of the five restoration strategies. Simulations used a landscape containing 1500 habitat cells (15% habitat cover), medium landscape heterogeneity, high within-patch habitat-quality heterogeneity, replicate 1, and an effective habitat area increase of 500 under the primary restoration-effort standardisation. At the start of each stochastic replicate, occupancy was assigned with probability $P_{init}=0.5$ only to cells available as habitat in the pre-restoration baseline landscape. The same initial occupancy realization and underlying stochastic random draws were used for the baseline and all restored states; consequently, newly restored habitat began unoccupied and had to be reached through colonisation.

Columns correspond to poor, intermediate, and good dispersers, corresponding to the lowest, middle, and highest representative values of the dispersal grid used in the main analysis ($\alpha=1$, $\alpha\approx21.5$, and $\alpha=1000$). Rows correspond to generalist, intermediate, and specialist species, corresponding to lowest, middle, and highest values of the specialisation grid ($\gamma=0.1$, $\gamma\approx0.77$, and $\gamma=10$). Each curve is based on 100 paired stochastic replicates at extinction-pressure parameter $e=0.5$.

The simulations qualitatively reproduced the principal trait-dependent differences observed in the metapopulation-capacity analysis. Improving habitat quality generally shifted persistence curves towards lower colonisation requirements for intermediate and specialist species, with particularly strong benefits for specialists, whereas improving matrix quality produced the strongest persistence benefits for generalists. Higher dispersal ability did not universally increase persistence in the SPOM because the dispersal kernel was normalised to sum to one; increasing $\alpha$ therefore redistributed colonisation pressure over greater distances rather than increasing its total magnitude. For specialists, whose persistence depends disproportionately on high-quality cells, this redistribution can reduce the concentration of colonisation pressure around the most suitable habitat. Overall, the stochastic simulations support the qualitative conclusion that the relative benefits of restoration strategies depend strongly on species traits, with habitat-quality improvement favoured increasingly with specialisation and matrix improvement particularly beneficial for generalists.
